# Offline hippocampal reactivation during dentate spikes supports flexible memory

**DOI:** 10.1101/2023.10.24.563714

**Authors:** Stephen B. McHugh, Vítor Lopes-dos-Santos, Manfredi Castelli, Giuseppe P. Gava, Sophie E. Thompson, Shu K.E. Tam, Katja Hartwich, David Dupret

## Abstract

Stabilizing new memories requires coordinated neuronal spiking activity during sleep. Hippocampal sharp-wave ripples (SWRs) in the Cornu Ammonis (CA) region, and dentate spikes (DSs) in the dentate gyrus (DG) are prime candidate network events supporting this offline process. SWRs have been studied extensively but the contribution of DSs remains unclear. By combining triple-(DG-CA3-CA1) ensemble recordings and closed-loop optogenetics in mice we show that, like SWRs, DSs synchronize spiking across DG and CA principal cells to reactivate population-level patterns of neuronal coactivity expressed during prior waking experience. Notably, the population coactivity structure in DSs is more diverse and higher-dimensional than that seen during SWRs. Importantly, suppressing DG granule cell spiking selectively during DSs impairs subsequent flexible memory performance in a multi-object recognition task and associated hippocampal patterns of neuronal coactivity. We conclude that DSs constitute a second offline network event central to hippocampal population dynamics serving memory-guided behavior.

## Introduction

Memories are stabilized during periods of sleep and rest (Maquet, 2001; Walker and Stickgold, 2006; Klinzing, Niethard and Born, 2019; Brodt *et al*., 2023). Decades of work have provided important insights into the underlying brain network mechanisms and have identified offline hippocampal activity as essential for this process (Buzsáki, 2010; Girardeau and Lopes-dos-Santos, 2021). Central to our current understanding are hippocampal sharp-wave ripples (SWRs): an intermittent, high-frequency (100 – 250 Hz) network event detected in the local field potentials (LFPs) of the CA1 region (Buzsáki, 1986, 2015; Axmacher, Elger and Fell, 2008; Joo and Frank, 2018). During SWRs, the firing activity of CA1 principal cells is transiently modulated (Csicsvari, H. Hirase, *et al*., 1999; Csicsvari *et al*., 2000) and reactivates the population-level firing patterns expressed in previous waking experience (Wilson and McNaughton, 1994). These offline spiking correlates have behavioral significance: suppressing CA1 neurons during SWRs impairs memory recall for recently acquired information (Girardeau *et al*., 2009; Ego-Stengel and Wilson, 2010; van de Ven *et al*., 2016). Conversely, prolonging SWRs or reinforcing the coordination between SWRs and neocortical activity promotes memory consolidation and subsequent behavioral performance (Maingret *et al*., 2016; Fernández-Ruiz *et al*., 2019). Hippocampal SWRs therefore constitute an offline network event important for memory-guided behavior. However, during sleep/rest periods the hippocampus exhibits another prominent network event: dentate spikes (DS), which are seen in the LFPs of the dentate gyrus (DG). To date, DSs have received little attention compared to SWRs. Accordingly, here we characterize the neuronal spiking dynamics nested in DSs with respect to SWRs and evaluate whether DSs constitute a second network event central to offline reactivation of waking firing patterns and subsequent memory-guided behavior.

The DG gates sensory information to the hippocampus, notably decorrelating these inputs into orthogonal patterns. This function may be crucial for the hippocampus to integrate multiple items in memory and to flexibly distinguish between stimuli with overlapping features (Treves and Rolls, 1994; McHugh *et al*., 2007; Knierim and Neunuebel, 2016). Within the DG, DSs represent intermittent, large amplitude network events recorded in the LFPs of the DG granule cell layer and are associated with increased spiking activity in dentate cells (Bragin *et al*., 1995; Senzai and Buzsáki, 2017; Dvorak *et al*., 2021). However, across the literature both increased and suppressed spiking activity of CA principal cells have been reported (Bragin *et al*., 1995; Penttonen *et al*., 1997; Meier *et al*., 2020; Dvorak *et al*., 2021; Sanchez-Aguilera *et al*., 2021), although, notably, some of these studies were in anesthetized (Penttonen *et al*., 1997; Sanchez-Aguilera *et al*., 2021) or head-fixed animals (Dvorak *et al*., 2021). Thus, here we further performed a systematic comparative assessment of DG and CA principal cell spiking activity during DSs versus SWRs in non-anesthetized, freely behaving mice.

To investigate the influence of DSs on hippocampal network activity and memory, we combined triple-(DG-CA3-CA1) site extracellular multichannel recordings and closed-loop optogenetic interventions in mice during active behavior and sleep/rest. We observed that DSs transiently increase principal cell spiking across the DG and CA regions of the hippocampus, nesting offline population-level activity patterns that are distinct from those in SWRs. Further, we report that DSs reactivate population activity patterns seen during prior waking experience. This DS-nested activity is relevant to whole-hippocampus population dynamics and memory-guided behavior: suppressing offline DG granule cell spiking selectively during DSs affects downstream CA principal cell spiking and impairs subsequent flexible memory performance in a hippocampal-dependent, multi-object recognition task. We propose that DSs constitute a second hippocampus network event that plays a complementary role to that of SWRs by supporting the offline reactivation of higher dimensional population patterns of neuronal coactivity in support of memory stabilization.

## Results

### Firing activity of hippocampal neurons synchronizes during DS events

We first used triple-(DG-CA3-CA1) site tetrode recordings to monitor network events in the LFPs and the spike trains of neuronal ensembles from the dorsal hippocampus of mice during sleep/rest (**Figure 1A,B**; n = 12 mice). From these LFPs, we detected DSs in DG and SWRs in CA1 to compare the spiking of principal cells between these two types of network event. Both DG DSs and CA1 SWRs were of short duration (median (IQR) duration: DSs = 42.4 (40.0 – 46.4) ms; SWRs = 47.5 (45.2 – 51.0) ms), occurring intermittently (median (IQR) occurrence frequency: DSs = 0.40 (0.25 – 0.58) Hz; SWRs = 0.75 (0.26 – 1.26) Hz) and rarely simultaneously (median (IQR) DS-SWR co-occurrence frequency: 0.03 (0.02 – 0.05) Hz). DS waveforms were consistent across mice and recording sessions (**Figure S1A,B**). We computed the firing responses of individual principal cells (n = 2,196 total recorded principal cells; CA1, 887; CA3, 388; DG, 921 cells) with respect to the peak of either DSs (without co-occurring SWRs) or SWRs (without co-occurring DSs). In line with previous work, DG principal cells transiently increased their firing activity during DSs (Bragin *et al*., 1995; Penttonen *et al*., 1997); the activity of CA principal cells increased during SWRs (**Figures 1C-G** and **S1C-H**) (Csicsvari, Hajime Hirase, *et al*., 1999; Buzsáki, 2015). We further observed that DG principal cell firing increased during SWRs (**Figures 1C,F,G** and **S1C-F**). However, we found that CA principal cells also increased their firing rate during DSs (**Figures 1D-G** and **S1C,G-H**), in contrast to some earlier reports that CA principal cell firing is suppressed during DSs (Bragin *et al*., 1995; Penttonen *et al*., 1997).

**Figure 1:**
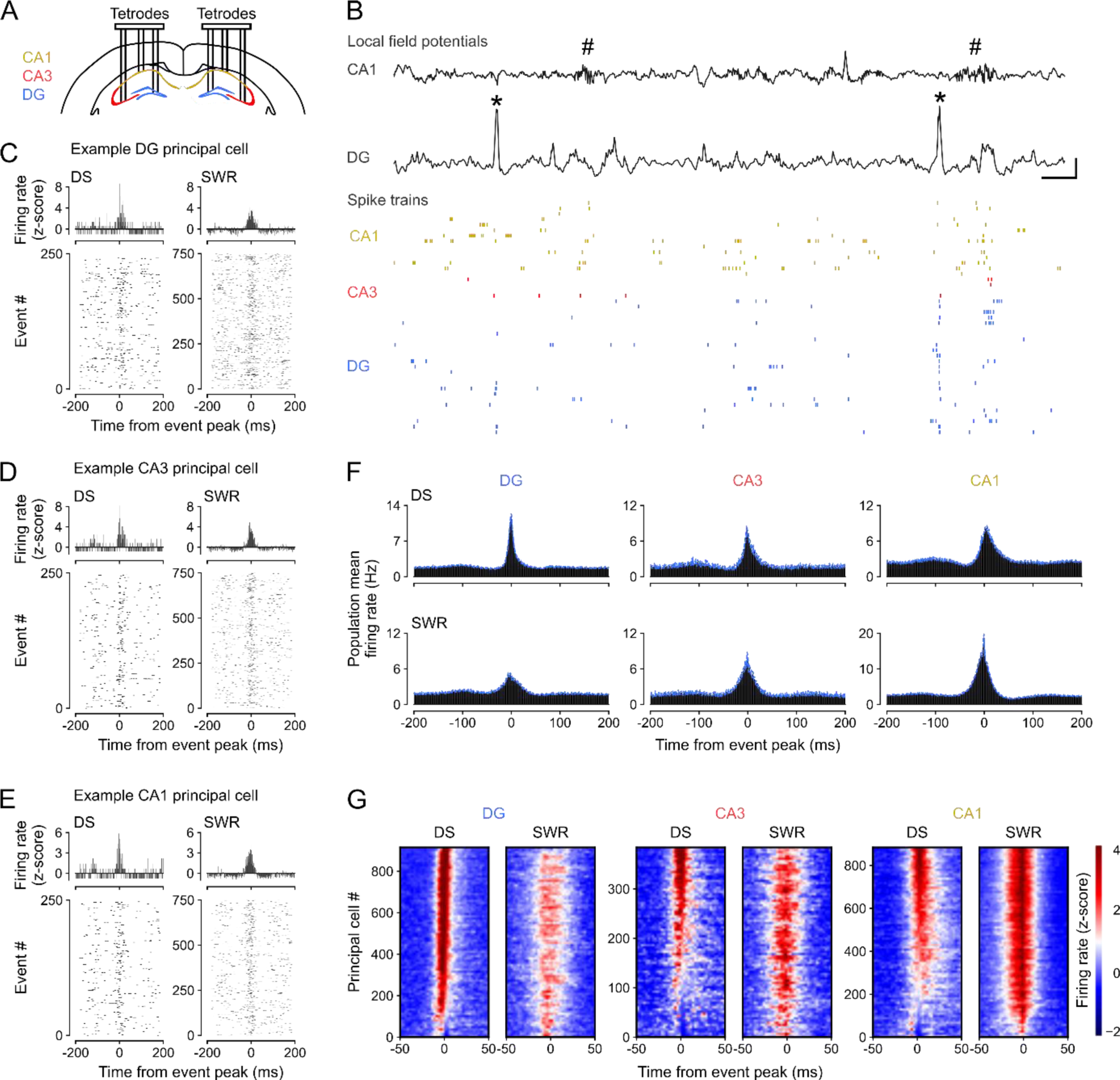
Dentate spikes recruit principal cell spiking across DG, CA3, and CA1. **(A)** Triple-(DG-CA3-CA1) ensemble tetrode recording allowed simultaneous monitoring of local field potentials (LFPs) and spiking activities. **(B)** Upper: raw wide-band CA1 and DG LFP traces (black) showing sharp-wave ripples (SWRs, hash symbols) in CA1 and dentate spikes (DSs, asterisks) in DG. Scale bars, 100 ms (horizontal), 1.5 mV for DG and 0.5 mV for CA1 (vertical). Lower: (color-coded) raster-plot of spike trains from CA1 (orange), CA3 (red), and DG (blue) principal cells (PCs, one cell per row). Shown is a few second sample of recording for clarity. **(C-E)** Spiking responses from single example DG (C), CA3 (D), and CA1 (E) principal cells. Upper: Z-scored peri-event time histogram (PETH) during DSs (left) and SWRs (right). Lower: corresponding raster plot showing event-related spiking responses (one event per row). **(F)** Group averaged firing rate PETHs for hippocampal PCs during DSs (top) and SWRs (bottom): DG (n=921), CA3 (n=388), CA1 (n=887) cells from 12 mice. Blue traces: mean ± SEM. **(G)** Heatmaps showing z-scored firing rates for the DG, CA3, and CA1 PCs shown in (F). For each heatmap: one cell per row, sorted (top-to-bottom) from the most activated (highest z-score at event peak, 0 ms, red) to the least activated (lowest z-score at event peak, blue) during DSs.

Previous studies identified two types of DS event (DS_1_ and DS_2_), based on the laminar profile of the transmembrane currents associated with the LFP expression of these network events (Bragin *et al*., 1995; Meier *et al*., 2020; Dvorak *et al*., 2021). Therefore, we next asked whether principal cell firing responses differed between DS_1_ and DS_2_. However, localizing sinks and sources of currents across hippocampal layers requires applying Current Source Density (CSD) analysis (Pettersen *et al*., 2012) to the LFPs measured at evenly spaced sites from the CA1 oriens layer to the DG granule cell layer. Such a laminar profile is not accessible with tetrode recordings. To distinguish between DS_1_ and DS_2_ events, we therefore implanted linear silicon-probes spanning the somato-dendritic axis of CA1 principal cells and reaching the inferior blade of the DG in a separate group of mice (n = 3). Having performed silicon-probe recordings during sleep/rest, we applied CSD analysis to these LFPs measured over the radial extent of the hippocampus to identify DS_1_ versus DS_2_ according to their underlying profile of current sinks and sources (**Figures 2A** and **S2A**; see methods). These CSD-validated DS_1_ and DS_2_ events exhibited distinct DG granule cell layer LFP waveforms (**Figures 2B** and **S2B**). We then trained a linear discriminant analysis classifier to identify these CSD-validated DS_1_ versus DS_2_ events using only their DG granule cell layer LFP signal. When tested on the silicon-probe LFP dataset, the classifier achieved over 85% accuracy (**Figure 2C**). When next applied to the tetrode LFP dataset, the classifier-identified DS_1_ and DS_2_ events also exhibited distinct granule cell layer LFP waveforms (**Figures 2D** and **S2C,D**), which were consistent with those obtained in silicon-probe recordings (**Figures 2B** and **S2B**). In both (tetrode and silicon-probe) datasets, DS_2_ represented two-thirds of all DS events (median (IQR): 66 (61 – 73) %), thus constituting the dominant type. Leveraging this cross-dataset approach, we found that the firing response of DG and CA principal cells was consistently stronger for DS_2_ than DS_1_ (**Figures 2E,F** and **S3A,B**). The average activity of principal cells in DS_1_ (and DS_2_) was nevertheless significantly higher than their baseline firing (calculated outside of any DS or SWR events during sleep/rest (**Figure S3C-F**). A greater proportion of CA principal cells showed suppressed firing activity during DS_1_ compared to DS_2_ (35% versus 11%; **Figures 2F** and **S3G,H**), providing insights into the previously documented DS-suppressed firing in some CA principal cells (Bragin *et al*., 1995; Penttonen *et al*., 1997; Nokia and Penttonen, 2022). Nevertheless, the proportion of active principal cells increased in DS_1_, DS_2_, and SWR events compared to baseline periods (**Figure 2G**), with equivalent proportions of neuronal recruitment in DS_2_ and SWRs (**Figure 2H**). These results show that the spiking activity of individual principal cells distributed across hippocampal regions is transiently modulated during DS events in offline sleep/rest behavior.

**Figure 2:**
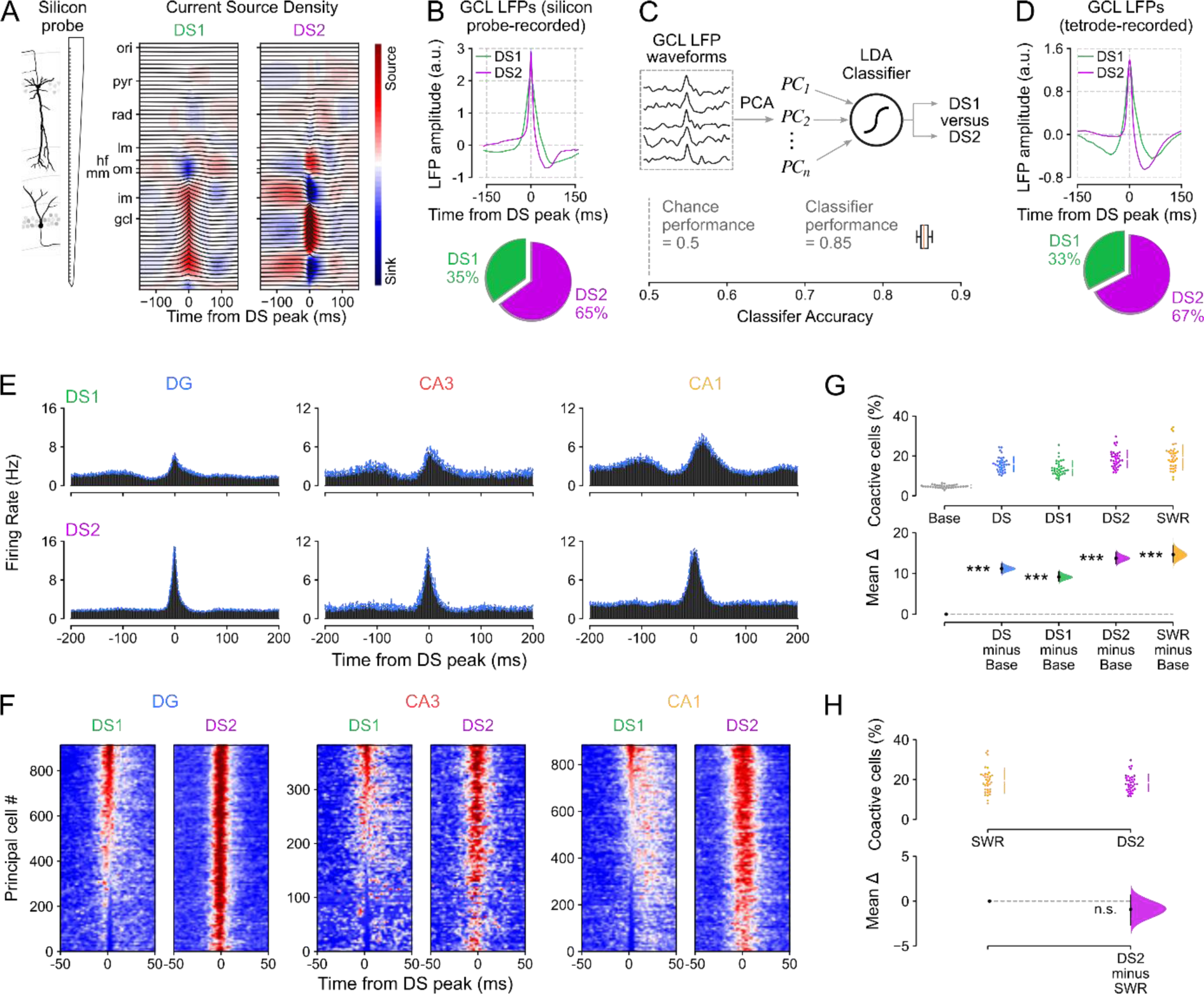
DS_2_ recruit more principal cell spiking than DS_1_ but similar levels to SWRs. **(A)** Left: Laminar (64-channel) silicon-probe recording allowed simultaneous monitoring of LFPs across hippocampal layers for current source density (CSD) analysis. Right: Example (radially organized) mean LFP traces (gray) with superimposed CSD profile (heatmaps) for type 1 (DS_1_) and type 2 (DS_2_) dentate spikes (calculated from 2,231 DS events in one mouse). Note the distinct CSD profiles reflecting the different transmembrane currents associated with DS_1_ versus DS_2_ events. Hippocampal layers: oriens (*ori*); pyramidale (*pyr*); radiatum (*rad*); lacunosum-moleculare (*lm*); outer (*om*), middle (*mm*), and inner (*im*) moleculare; granulare (*gcl*). Hippocampal fissure (*hf*). **(B)** Upper: Shown for silicon-probe recorded DS_1_ and DS_2_ identified from their CSD profiles are example average granule cell layer LFP waveforms triggered by the peak of these events. Lower: these recordings showed a higher proportion of DS2 than DS1 events (n=15,067 events, 3 mice). **(C)** Upper: we applied principal component analysis on the normalized granule cell layer LFP waveforms for all silicon-probe recorded DS events. We then used the principal components explaining 90% of the variance to train a linear discriminant classifier with the true labels (DS_1_ versus DS_2_) determined by the individual CSD profiles. Lower: the classifier performance (>85%) was significantly above chance level (50%) when tested on silicon-probe recorded LFP waveforms of unlabeled events. We used this classifier to next distinguish DS_1_ and DS_2_ from tetrode-recorded granule cell layer LFP waveforms (D). **(D)** Upper: Shown for tetrode-recorded DS events are the average granule cell layer LFP waveforms for DS_1_ and DS_2_ predicted label obtained from the silicon-probe-based classifier (C). Lower: these recordings also showed a higher proportion of DS_2_ than DS_1_ events (n=32,215 events, 12 mice). **(E)** Group averaged firing rate PETHs for tetrode-recorded DG, CA3, CA1 principal cells during DS_1_ and DS_2_ (as Figure 1F,G). Blue traces: mean ± SEM. **(F)** Heatmaps showing z-scored firing rates for the DG, CA3, and CA1 cells shown in (E). For each heatmap: one cell per row, sorted (top-to-bottom) from the most activated (highest z-score) to least activated (lowest z-score) during DS_1_ peaks. **(G)** Estimation plot showing the effect size for the differences in the percentage of cells coactivated during SWRs and DSs (with DS_1_ and DS_2_ plotted altogether or separately) and compared to equivalent (50 ms duration matched) baseline windows (Base) without any co- occurring DS events. Upper: raw data points (each point represents one session with at least 100 of each event type and 20 principal cells), with the gapped lines on the right as mean (gap) ± s.d. (vertical ends) for each event. Lower: difference (Δ) in percentage of cells coactivated between Base windows and all DS, DS_1_, DS_2_, and SWR events computed from 5,000 bootstrapped resamples and with the difference-axis origin (dashed line) aligned to the baseline percentage of coactive principal cells (black dot, mean; black ticks, 95% confidence interval; filled curve, sampling-error distribution). **(H)** As (G) but comparing percentage of coactive cells during SWR versus DS_2_. Note that DS_2_ and SWR events engage similar levels of neuronal activity. \*\*\**P* < 0.001.

### DS events nest higher dimensional patterns of population coactivity

We next investigated how the hippocampus organizes the collective activity of its principal cells both within individual DS events and across events, comparing these population-level patterns to those expressed in SWRs. To proceed, we first considered the neuron-wise vectors formed by the instantaneous spike discharge of principal cells in DSs, SWRs, or duration-matched control windows (without any DSs or SWRs) of the same sleep/rest (**Figures 3A** and **S4A**; “population vector analysis”). A logistic regression classifier trained on these population firing vectors significantly distinguished DS from SWR events but could not distinguish between their corresponding pre-event control epochs (**Figure 3B**). Successful classification was also obtained when using only DS_2_ events, which matched SWRs in the proportion of cells activated per event (**Figure S4B**). When evaluating the pairwise similarity of DS-nested population vectors versus those of SWR-nested vectors (**Figure 3C**), we further observed that DSs contained a higher diversity of firing vectors compared to those in SWRs, which were more similar to one another (**Figures 3D** and **S4C,D**).

**Figure 3:**
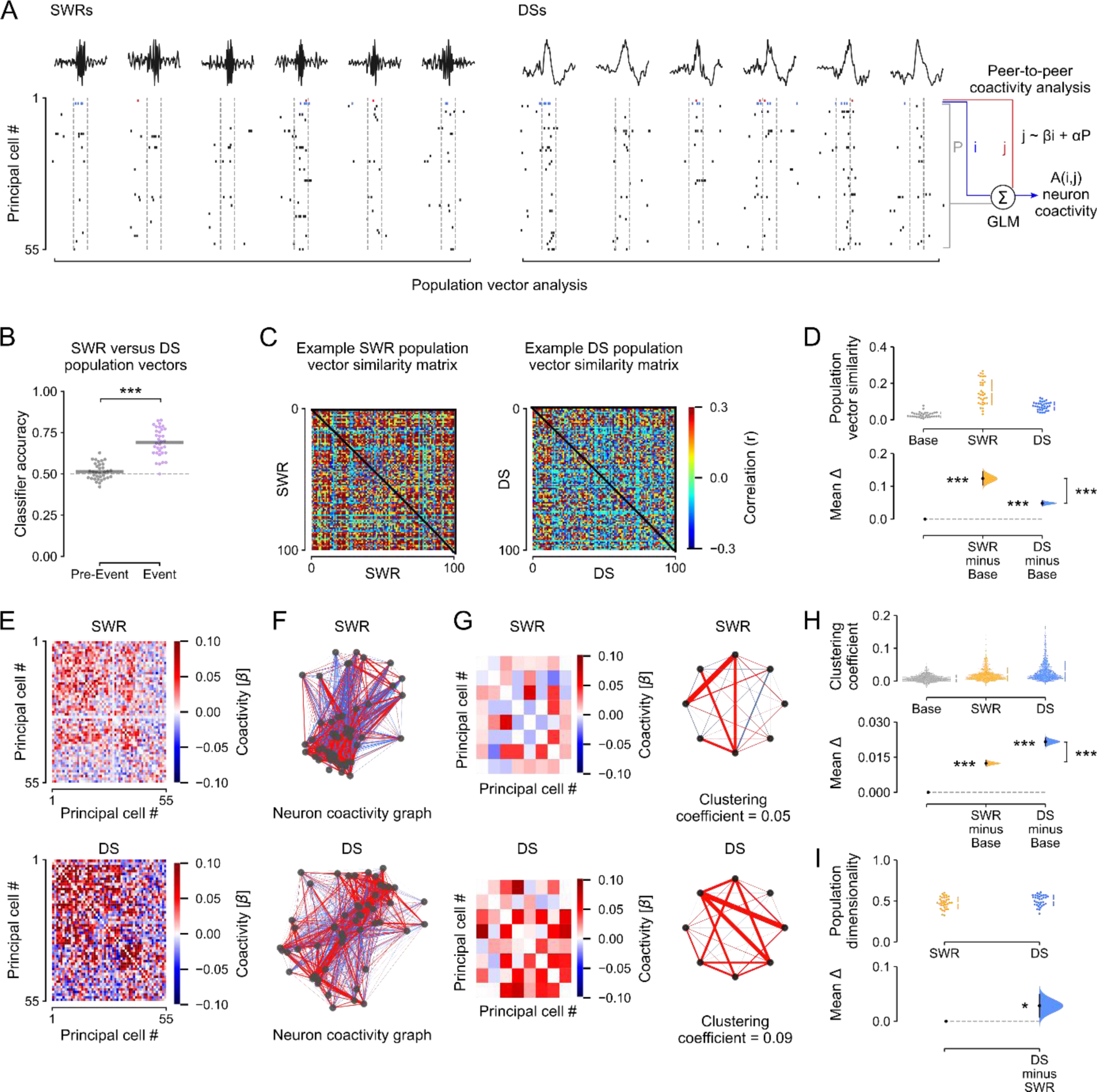
The coactivity structure of population spiking differs between DSs and SWRs. **(A)** Analytical framework: the population-level coactivity structure was analyzed using population vectors of principal cell spiking transiently nested in individual SWRs, DSs, or duration-matched (50 ms) control windows. For the analyses in panels B-D, these population firing vectors were then binarized (for each cell: a non-zero spike count gives 1; or else 0). For the analyses in E-H, we calculated the peer-to-peer coactivity, controlling for the overall population activity. **(B)** A logistic regression classifier trained on population vectors nested in SWR versus DS events, or matched duration pre-event control windows, using a 4-fold cross-validation approach (75% of vectors for training; the remaining 25% for evaluation), significantly discriminated DSs from SWRs, but could not discriminate between pre-DS versus pre-SWR vectors. Gray horizontal bars: mean classification accuracy. **(C-D)** DS population firing vectors are more diverse than those in SWRs. For each sleep session, we computed the similarity (Pearson correlation coefficient) for each pair of population vectors nested in either DSs, SWRs, or duration-matched baseline windows (Base). (C) shows example DS and SWR matrices of cross-vector similarity for one session. Cross-population vector similarity was significantly higher in SWRs compared to both DSs and control windows (D). **(E-H)** DS and SWR population firing vectors exhibit distinct topology of neuronal coactivity. The coactivity between any two (*i*, *j*) neurons was measured using a GLM that quantified their short timescale (50 ms windows centered on DS or SWR peaks) firing relationship while accounting for network-level modulation reported by the remaining principal cells in the population (A). (E) This procedure returned for both DS and SWR events an adjacency matrix of *β* regression weights that represented the neuron pairwise coactivity structure of the population. (F) Visualization of the corresponding matrices representing DS and SWR based neuronal coactivity graphs. For clarity, (G) shows an example subset (left) for each adjacency matrices shown in (E), along with its corresponding motifs of neuronal coactivity and clustering coefficient (right). (H) Note that DS-based graphs contained more triads of coactive nodes compared to both SWR graphs and control graphs constructed from duration-matched baseline windows (Base), as indicated by higher mean clustering coefficients **(I)** The dimensionality of population vector matrices (number of principal components required to explain 90% of the variance) was higher for DSs than SWRs. \**P* < 0.05, \*\**P* < 0.01, \*\*\**P* < 0.001.

This difference in cross-population vector similarity suggested that DSs and SWRs differ with respect to their neuronal motifs of transient coactivation. By examining the topological organization of peer-to-peer firing associations, we indeed observed that DS events contain more motifs of coactive principal cells than SWRs. For each cell pair (*i*, *j*), we trained a generalized linear model to predict the spike discharge of neuron *j* from that of neuron *i* while accounting for the activity of the remaining peers (**Figure 3A**; “peer-to-peer coactivity analysis”). We performed this procedure separately for DS and SWR events, which returned for each type of network event a matrix of *β* regression weights that represented the coactivity structure of the population (**Figure 3E**). With these matrices, for both DS and SWR events we constructed neuronal coactivity graphs (with no self-connections) where each node is a cell and the edge linking any two nodes represents the firing association of that cell pair (**Figure 3F,G**). This revealed that DS-based graphs contained more triads of coactive nodes compared to SWR graphs (**Figure 3H**). This remained the case when comparing DS_2_ and SWR events (**Figure S4E**).

These findings showed that while both DS and SWR events synchronize hippocampal principal cells, population coactivity responses to DSs are more diverse. The higher diversity of DS firing vectors could allow the hippocampus network to transiently expand to higher- dimensional activity subspaces where more population patterns can coexist with minimized interference. Consistent with this idea, applying principal component analysis showed higher dimensionality of population vectors in DS events compared to SWRs (**Figures 3I** and **S4F-G**).

### Waking theta coactivity patterns reactivate in offline DSs and support flexible memory

The DS-nested motifs of peer-to-peer firing associations could instantiate population patterns of neuronal coactivity undergoing offline reactivation to support memory-guided behavior.

Notably, the link between hippocampal SWRs and memory reactivation was initially established through the observation that the neural patterns of joint spiking activity expressed during exploratory behavior are more strongly correlated with those nested in post-exploration sleep/rest SWRs than those in SWRs before waking experience (Wilson and McNaughton, 1994; O’Neill *et al*., 2008). Accordingly, we next determined whether DSs constitute another hippocampal timeframe for offline reactivation of waking coactivity patterns. To proceed, we used our peer- to-peer coactivity analysis (**Figure 3A**), applying it to DS versus SWR events of sleep/rest before and after exploration of open field arenas (**Figure 4A**). Likewise, we obtained the waking patterns of population coactivity in theta cycles during exploration (**Figure 4A**). With these, we computed DS and SWR reactivation by measuring the tendency of the peer-to-peer theta firing associations to reoccur in post-exploration sleep/rest DS (or SWR) events, while controlling for prior pre-exploration DS (or SWR) coactivity. In line with previous work, offline patterns of SWR coactivity reflected those of theta coactivity significantly more during post-exploration than pre-exploration sleep/rest (**Figure 4B**; *left panel*). This SWR reactivation was significantly higher than that obtained with a null distribution generated from models using randomly shuffled cell pair identities (**Figure 4B**; *right panel*). Importantly, we observed that theta coactivity patterns were also strongly reactivated in post-exploration DSs (**Figure 4C** and **Table S1**). By applying the same analytical framework to the neuronal ensembles tracked in both SWRs and DSs, this result provided evidence for offline DS reactivation of hippocampal waking firing patterns.

**Figure 4:**
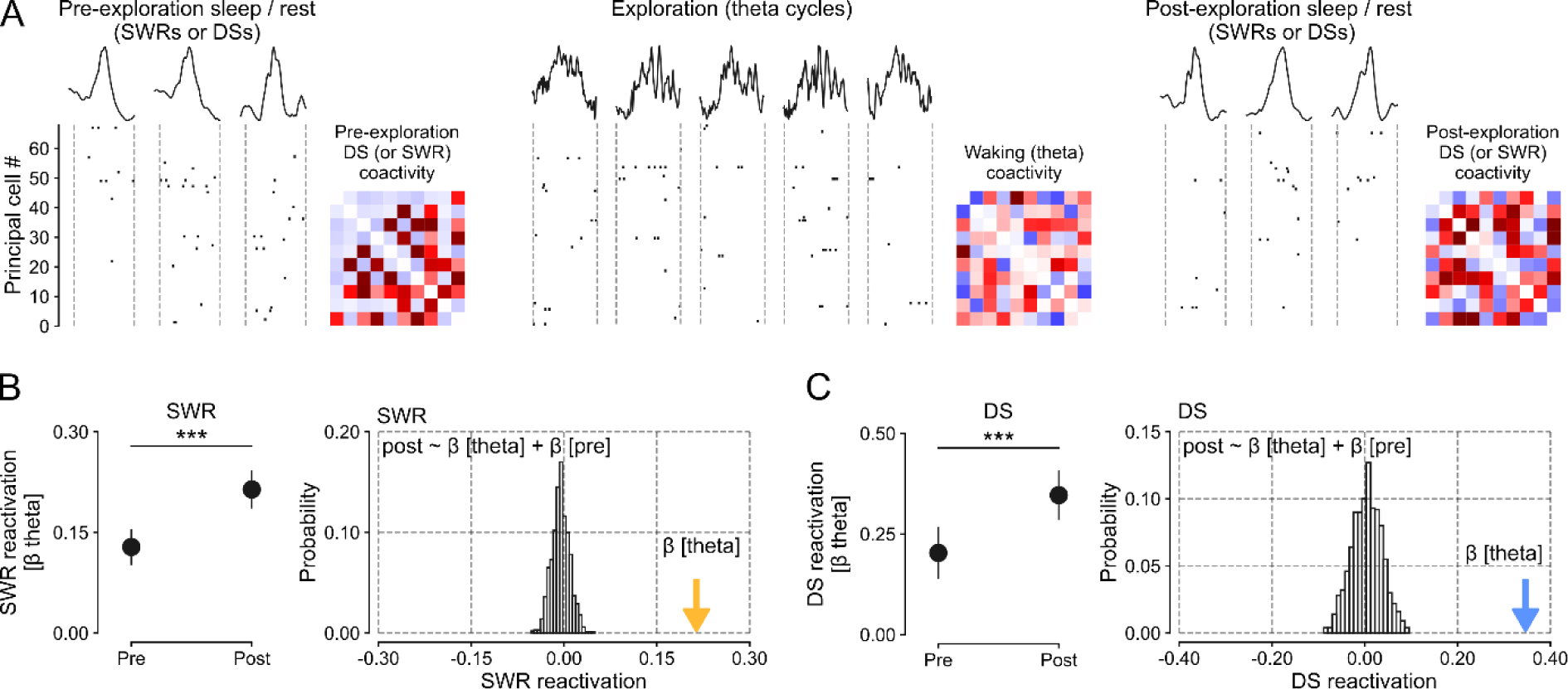
Waking patterns of hippocampal coactivity reactivate in offline DSs. **(A)** DS and SWR reactivation of waking patterns formed by principal cell theta coactivity. We compared the tendency of principal cell pairs to co-fire in theta cycles during exploration (waking theta coactivity) with their tendency to co-fire in DS or SWR events during the following sleep/rest period (post-exploration DS or SWR coactivity), controlling for their baseline co-firing in the sleep/rest period before (pre-exploration DS or SWR coactivity). Inset matrices show example ‘peer-to-peer’ coactivity matrices for each behavioral stage. **(B)** SWR reactivation (measured by the β coefficients of the linear regression that predicted post- exploration SWR coactivity from waking theta coactivity, controlling for pre-exploration SWR coactivity). Left: The β coefficient for theta coactivity was significantly higher when predicting post-exploration SWR coactivity than with the reverse model (i.e., predicting pre-exploration SWR coactivity from theta coactivity, controlling for post-exploration SWR coactivity). Error bars show ± 95% confidence interval. Right: The histogram shows the random probability distribution of β weights for theta coactivity when cell pair identity was shuffled (i.e., a null distribution based on 1,000 random shuffles; n=7,010 cell pairs). The colored arrow shows the actual β coefficient for theta coactivity. **(C)** DS reactivation exhibited the same pattern of results as SWR reactivation, shown in B. \*\*\**P* < 0.001.

The reactivation of waking population patterns in DSs (**Figure 4C**), which contain more diverse and higher-dimensional patterns of neuronal coactivation (**Figure 3D,H,I**), raised the question of their network contribution to memory-guided behavior. We thus tested whether the offline population response during DSs was necessary for a task that requires integrating multiple items in memory, to flexibly distinguish between previously-encountered versus novel objects. To this end, we transduced dentate granule cells with the yellow (561-nm) light-driven optogenetic silencer Archaerhodopsin T (ArchT) in Grm2-Cre mice (**Figure 5A,B**). We subsequently implanted these Grm2^DG^::ArchT mice for triple-(DG-CA3-CA1) ensemble recordings combined with bilateral optic fibers for DG light delivery. In these experiments, DG light delivery was performed in a closed-loop manner during sleep using real time monitoring of DS onset (**Figure 5A**; “DS-sync” condition). We used a within-subject control paradigm where, on different days, light was instead delivered after each DS had elapsed (“DS-delay” condition). Light delivery synchronized to DS onset significantly suppressed principal cell activity compared to the delay condition (**Figures 5C** and **S5A,B**). We applied this approach during interposed sleep sessions in a hippocampal-dependent multi-object recognition memory task (McHugh *et al*., 2022). In this task, mice repeatedly explored a square-walled arena containing four objects (**Figure 5D**). In the first session (‘Sampling’), mice encountered four distinct novel objects, each one placed beside a wall. On the subsequent sessions (’Test’), one of the initially sampled objects was replaced by a different novel object so that the mouse could explore one completely novel object along with the three ‘familiar’ objects from the previous session that day. These exploration (sampling and test) sessions alternated with sleep/rest sessions where mice received DG-targeted light delivery, either synchronized or delayed with respect to DS onset. In each test session *n*, we measured novelty preference using the proportion of time mice spent investigating the novel versus the familiar objects, thereby probing recognition memory for objects explored in session *n* − 1. We found that novelty detection was not impaired in test sessions following sleep with DS-delayed suppression, with mice expressing a stronger preference for novel over familiar objects (**Figure 5E**). This novel object preference was similar to that observed in control mice without any optogenetic intervention (**Figure S5C**). However, novel object preference was absent in test sessions following DS-synchronized suppression (**Figures 5E** and **S5D**). The total object exploration time did not differ between the DS- synchronized and DS-delayed conditions (**Figure S5E**). We also observed that successful recognition memory in the DS-delayed condition was accompanied by stronger peer-to-peer theta coactivity during exploration of familiar compared to novel objects (**Figure 5F**). This difference in theta coactivity was not present following DS-synchronized hippocampal suppression (**Figure 5F**). Moreover, in the DS-delayed condition, theta coactivity increased over test sessions whereas it decreased in the DS-synchronized condition (**Figure S5F**). Thus, the hippocampal population response to offline DS events is required for flexible, memory-based recognition of previously encountered items and associated network increase in theta coactivity patterns.

**Figure 5:**
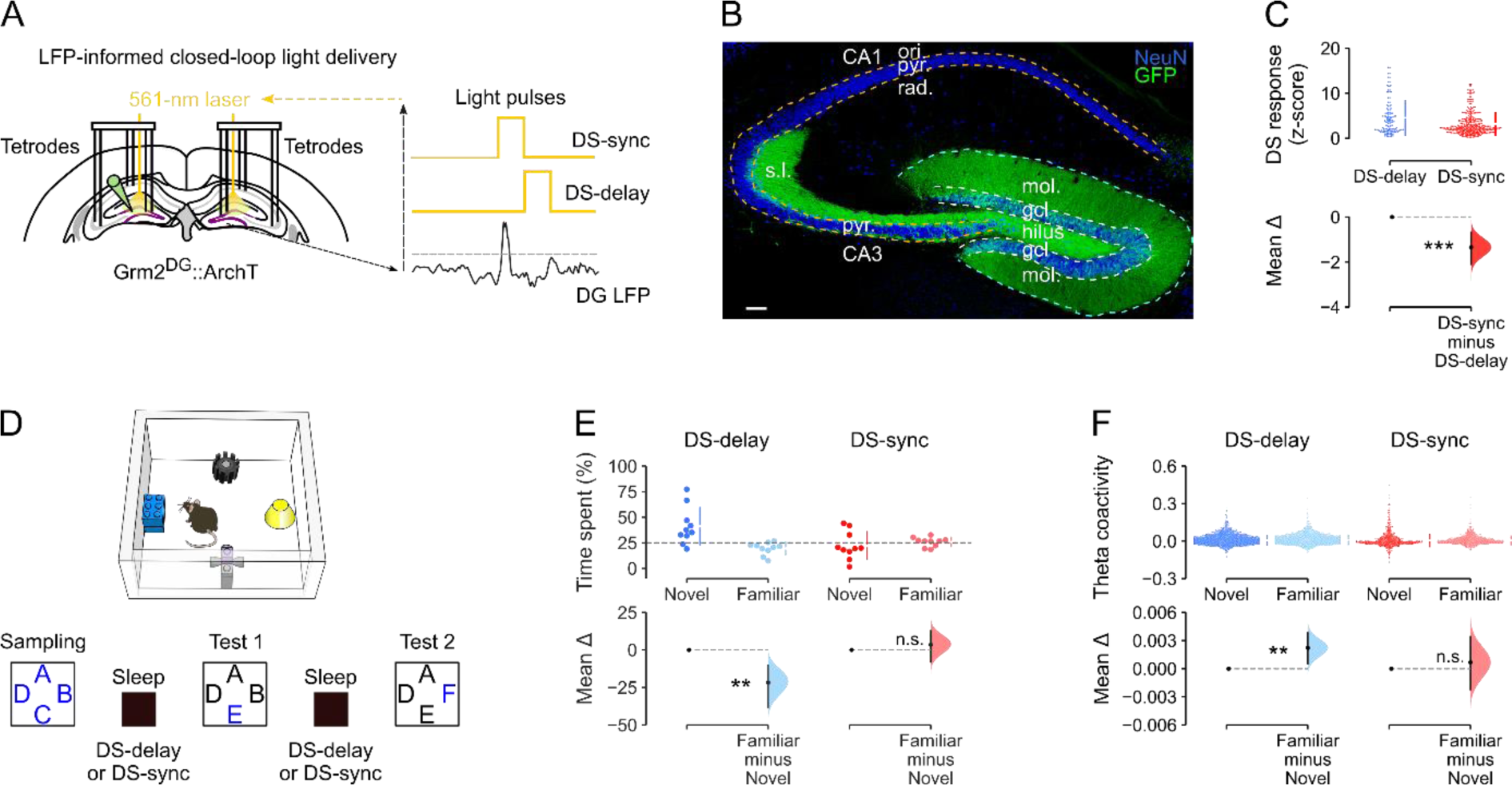
Offline suppression of DS activity impairs subsequent recognition memory. **(A)** Triple-(DG-CA3-CA1) ensemble recording with LFP-informed yellow (561 nm) DG light- delivery. Dentate granule cells (DGCs) transduced with ArchT-GFP (Grm2^DG^::ArchT). Closed- loop light-delivery to suppress DGC spiking immediately upon DS-detection (DS-sync condition) and the control condition (DS-delay, where light delivery was offset by 100 ms after DS detection). **(B)** ArchT-GFP-expressing DGCs in a Grm2^DG^::ArchT mouse. Neuronal nuclei stained with NeuN. Scale bar=100 µm. Granule cell layer: gcl; molecular layer: mol; pyramidal cell layer: pyr; stratum oriens: ori; radiatum: rad; lucidum: s.l. **(C)** Estimation plot showing the effect size for the difference in (z-scored) firing rate of DGCs during DS-delay (blue) versus DS-sync (red) conditions. Upper: each data point represents one cell. Lower: mean difference between the two conditions (DS-delay: n=127; DS-sync: n=319 DGCs in 5 mice). **(D)** Task arena (upper) and layout (lower). Letters depict object locations (novel objects in blue). During sleep sessions (interposed between novel object exploration sessions), mice received either DS-synchronised (DS-sync) or delayed (DS-delay) DGC suppression. **(E)** Estimation plot showing the percentage of time spent by mice with the novel versus the familiar objects in the DS-delay and DS-sync conditions. Upper: Each data point represents the mean time spent by one mouse with the novel object versus the three familiar objects; chance performance is shown by the dashed line. Lower: mean difference between novel and familiar object exploration time. Mice in the DS-delay condition, but not the DS-sync condition, exhibited a significant preference for novel over familiar objects. **(F)** Theta-paced peer-to-peer coactivity during object exploration was higher for familiar than novel objects in the DS-delay condition but not the DS-sync condition. \*\**P* < 0.01, \*\*\**P* < 0.001.

## Discussion

Our findings establish that offline DSs activate neurons across the DG and CA regions. DSs therefore constitute a second hippocampal network event that hosts short timescale coactivation forming population-level neural patterns, like the well-established SWRs. However, the structure and content of spiking activity is distinct in DSs. Notably, we found that DSs nest more motifs of coactive neurons, yielding population patterns of higher diversity and dimensionality compared to those in SWRs. Like SWRs, DSs reactivate hippocampal population patterns expressed in prior waking experience. This offline reactivation is behaviorally significant: closed-loop suppression of DG granule cell spiking selectively during offline DS events is sufficient to disrupt downstream CA principal cell activity and impair subsequent flexible recognition memory for previously encountered items, as well as the associated network gain in theta-nested neuronal coactivity. Collectively, these findings identify a core contribution for DSs to hippocampal patterns of population coactivity and memory-guided behavior.

We started this investigation by observing that DSs increase spiking activity in principal cells across the DG, CA3, and CA1 regions of the hippocampus. This finding is consistent with previous reports of DS-evoked spiking activity in DG granule cells but contrasts with some earlier reports of DS-suppressed CA pyramidal cell spiking (Bragin *et al*., 1995; Penttonen *et al*., 1997; Meier *et al*., 2020; Dvorak *et al*., 2021). Notably, Bragin and colleagues reported suppressed spiking in 3/14 CA3 principal cells and suppressed CA1 multi-unit activity in 2/10 rats, showing some, rather than consistent, CA suppression. In addition, Penttonen et al. reported suppressed CA1 multi-unit activity and hyperpolarization of 4 intracellularly recorded CA1 neurons during DSs in anaesthetized rats. However, DS rates are ∼10-fold lower and have smaller amplitude during anesthesia compared to DSs observed during natural sleep/rest (Penttonen *et al*., 1997). Other studies reported increased CA1 multi-unit activity (Meier *et al*., 2020) and increased CA3 single-unit spiking during DS events (Dvorak *et al*., 2021) in non- anaesthetized mice. Here, we report a variety of firing responses across individual CA neurons, ranging from strong activation to suppression during DS events (**Figures 1G** and **2F**). However, our systematic study (including > 2,000 principal cells) shows that DSs do drive increased mean population spiking activity in both DG and CA principal cells.

Previous studies distinguished between two types of DS event (DS_1_ and DS_2_), based on the laminar profile of their transmembrane currents (Bragin *et al*., 1995; Meier *et al*., 2020; Dvorak *et al*., 2021). Here, we found that DS_2_ are more effective than DS_1_ events at recruiting hippocampal principal cells, with higher spike rates per cell and more coactive cells per event. This result is consistent with a recent report that DS_2_ but not DS_1_ events reliably increase spiking in DG and CA3 principal cells (Dvorak *et al*., 2021). The same report saw only a slight increase in CA1 spiking during DS_2_, and no effect of DS_1_ events on either CA1 or CA3 principal cells.

They also found that DG principal cell spiking was suppressed during DS_1_, in contrast to our study and previous reports (Bragin *et al*., 1995). Here we found that both DS_1_ and DS_2_ events evoked significantly increased spiking activity in DG, CA3, and CA1 principal cells, but again would emphasize the diversity of CA cell responses, especially during DS_1_ events (**Figure 2F**). The observed differences between studies might reflect differences between DS events in head- fixed awake mice versus those in sleep/rest (Bragin *et al*., 1995; Headley, Kanta and Paré, 2017), akin to the differences previously reported for SWRs detected across distinct behavioral states, from exploration to quiet immobility to sleep (O’Neill, Senior and Csicsvari, 2006). These observed differences could also indicate a moment-to-moment diversity across DS events, similar to that recently highlighted for individual SWRs (Navas-Olive *et al*., 2022) and theta cycles (Lopes-dos-Santos *et al*., 2018).

In this study, we directly compared patterns of population spiking activity in DSs versus SWRs. Strikingly, we found that the structure of population activation differed across these two types of brief network event; allowing a classifier to distinguish DSs versus SWRs based on their population vectors of principal cell spiking. We also found that the tendency of groups of neurons to fire together was stronger during SWRs than DSs, in line with the observation of more (uncorrelated) activity patterns and therefore higher dimensionality in DSs (**Figure 3**).

Importantly, we found that waking patterns of neuronal coactivity nested in theta oscillations reactivate in DSs of post-exploration sleep/rest (**Figure 4**). These findings thus provide important evidence for offline reactivation of hippocampal waking firing patterns outside of SWRs.

To determine whether spiking activity observed during DS events was required for subsequent memory-guided behavior, we deployed a closed-loop optogenetic feedback approach to suppress DG granule cell activity selectively during DS events. This DS-synchronized suppression of DG principal cells reduced spiking activity in CA principal cells. When applied in sleep/rest following object exploration, this DS-synchronized neural suppression impaired subsequent novel object recognition memory performance. Although ours is the first study to leverage a closed-loop optogenetic approach, this finding is consistent with previous behavioral studies using electrical stimulation to disrupt hippocampal activity during DSs (Nokia *et al*., 2017; Lensu *et al*., 2019; Nokia and Penttonen, 2022). Moreover, we found that increased theta coactivity was associated with familiarity, both within and across test sessions; and that this gain in theta coactivity was absent following DS-synchronized neural suppression. This finding is consistent with recent work showing that continual integration of new memory items across behavioral experiences increases neuronal coactivity (Gava *et al*., 2021). Altogether, these results support the idea that neuronal activity during DS events plays an important role in subsequent memory-guided behavior, as SWRs do.

Why does the hippocampus use more than one offline network mechanism to support memory? DSs and SWRs are driven by distinct neural circuits. SWRs depend on excitatory inputs from CA3, generating high-frequency CA1 ripples (Buzsáki, 1986; Nakashiba *et al*., 2009; Sullivan *et al*., 2011; Davoudi and Foster, 2019). DSs are non-oscillatory events driven by excitatory inputs from the entorhinal cortex (Bragin *et al*., 1995; Headley, Kanta and Paré, 2017; Dvorak *et al*., 2021), lesions of which eliminate DSs but increase SWR incidence (Bragin *et al*., 1995). Our structural analysis of DS versus SWR population patterns raises the intriguing possibility that SWRs may be more suited for lower-dimensional network coactivity serving robust information flow; whereas DSs may promote higher-dimensional activity, allowing diverse mnemonic patterns to coexist offline and support flexible, pattern separation during subsequent behavior. Collectively, these findings open important new avenues for future work to explore the interplay between DS versus SWR events as two distinct timeframes for the hippocampus to optimize offline computations serving memory-guided behavior.

## Materials and Methods

### Animals

These experiments used adult (4–6 months old) wild-type C57Bl6/J mice (Charles River Laboratories, Kent, UK) or hemizygous Nestin-Cre mice (Jackson Laboratories; B6.Cg-Tg(Nes-Cre)1Kln/J, stock no. 003771, RRID: IMSR_JAX:003771) for the initial investigation of principal cell spiking in DSs and SWRs using tetrodes or silicon-probe recordings (**Figures 1–4**). To optogenetically target DG granule cells, we used adult metabotropic-glutamate-receptor 2- Cre (Grm2-Cre) hemizygous male mice (**Figure 5**). This Grm2-Cre mouse strain was obtained from the Mutant Mouse Resource and Research Center (MMRRC; Tg(Grm2- cre)MR90Gsat/Mmucd; stock no. 034611-UCD, RRID:MMRRC_034611-UCD) at University of California at Davis, an NIH-funded strain repository, and was donated to the MMRRC by Nathaniel Heintz, Ph.D., The Rockefeller University, GENSAT and Charles Gerfen, Ph.D., National Institutes of Health, National Institute of Mental Health. All mice were group housed with same-sex littermates until the start of the experiment and singly housed after surgery. Mice had free access to food and water throughout, in a dedicated housing room with a 12/12 h light/dark cycle (7 a.m.–7 p.m.), 19–23 °C ambient temperature and 40–70 % humidity. All experiments were performed between 8 a.m.–6 p.m. during the light-on period, that is when mice sleep more. Experiments were performed in accordance with the Animals (Scientific Procedures) Act, 1986 (United Kingdom), with final ethical review by the Animals in Science Regulation Unit of the UK Home Office.

### Viral vectors

An AAV carrying a double-floxed inverse open reading frame (DIO) Cre-dependent opsin under the CAG promoter was used to deliver Archaerhodopsin (ArchT) (Han *et al*., 2011) into DG granule cells (AAV9-CAG-Flex-ArchT-GFP, titer: 8.3 × 10^12^ TU / mL, University of North Carolina).

### Surgical procedures

Mice received viral injections and microdrive implantations under gaseous isoflurane anaesthesia (∼1% in 1 L / min O_2_), with systemic and local analgesia administered subcutaneously (meloxicam 5 mg / kg; buprenorphine 0.1 mg / kg; bupivacaine 2 mg / kg).

Viruses were injected bilaterally into the dorsal DG (3 × 200 nL per hemisphere; at the following stereotaxic coordinates from bregma: anterior-posterior: -1.6, -2.4, -2.4; mediolateral: ±1.0, ±1.2, ±1.5; dorsoventral: -1.7, -1.7, -1.7 mm, respectively), and delivered using a pulled glass micropipette (∼16-µm i.d.) at a rate of 100 nL min^−1^, with an additional 100 nL min^−1^ diffusion time with the pipette *in situ*. In a separate surgery, mice were implanted with a microdrive containing twelve- or fourteen-independently movable tetrodes bilaterally targeting DG, CA3 and CA1, and two optic fibers (Doric Lenses Inc., Quebec, Canada) positioned bilaterally above the dorsal DG. A separate group of mice were implanted with unilateral single-shank 64-channel silicon-probe (model: ASSY-236 H3, 8-mm; Cambridge Neurotech, Cambridge, UK; stereotaxic coordinates from bregma: anterior-posterior: -2.0; mediolateral: -1.7 mm); these mice did not receive prior viral injections.

### Recording procedures

Following the implantation surgery, mice recovered for at least seven days before familiarization to the recording procedure. Mice were handled daily and exposed to the sleep-box for > 0.5 h per day for at least four days. During this period, tetrodes / silicon-probes were slowly lowered to the proximity of the cell layers. Once at the correct depth, silicon-probes were left in the same position for the rest of the experiment. Tetrodes were lowered into the CA1, CA3 pyramidal or DG granule cell layers on the morning of each recording day in search of multi-unit spiking activity, using the electrophysiological profile of the local field potentials including sharp-wave ripples, gamma oscillations, and dentate spikes to further guide placement. Tetrodes were left in position for ∼1.5–2 h before recordings began on that day. At the end of each recording day, tetrodes were raised (∼150 µm) to protect the cell layer overnight.

During recording sessions, mice explored open-field environments (41-cm diameter cylinder, or 41 × 41-cm square box, both with 30-cm high walls), or were placed in a sleep box containing sawdust bedding and nesting material (12 × 12 × 28-cm, length × width × height). Each recording session lasted ∼15-30 min. Experiments were performed under dim light conditions (∼20 lux) with low-level background noise (∼50 dB).

### Light delivery

A 561-nm diode pumped solid-state laser (Crystal Laser, model CL561-100; distributer: Laser 2000, Ringstead, UK) was used to deliver green-yellow light bilaterally to the dorsal DG (∼5-9 mW) via a 2-channel rotary joint (Doric Lenses Inc.).

### Multichannel data acquisition

Electrode signals were amplified, multiplexed, and digitized using a single integrated circuit (headstage) located on the head of the animal (RHD2164, Intan Technologies, USA; http://intantech.com/products_RHD2000.html). The amplified and filtered (pass band 0.09 Hz to 7.60 kHz) electrophysiological signals were digitized at 20 kHz (RHD2000 Evaluation Board) and saved to disk with the synchronization signals from the positional tracking and laser activation. To track the location of the animal, three LEDs were attached to the headstage and captured at 25 frames per second by an overhead color camera.

### Spike sorting and unit isolation

Spike sorting and unit isolation were performed via automatic clustering software Kilosort (Pachitariu *et al*., 2016, 2023) (https://github.com/cortex-lab/KiloSort) followed by graphically based manual recombination using cross-channel spike waveforms, auto-correlation histograms and cross-correlation histograms within the SpikeForest framework (https://github.com/flatironinstitute/spikeforest) (Magland *et al*., 2020). All sessions recorded on a given day were concatenated and cluster cut together to monitor cells throughout the day. Units that were well-isolated and stable over the entire recording were used for analysis. Hippocampal principal cells were distinguished from interneurons by their auto-correlograms, firing rates, and spike waveforms as described previously (Csicsvari *et al*., 1999). In total, this study includes 2,777 hippocampal principal cells (CA1, n = 1,139; CA3, n = 452; DG, n = 1,186; from 103 total recording days in 21 mice).

### Local field potential signals

LFP signals were processed by first applying an anti-aliasing filter (8^th^-order Chebyshev type I filter) to the wide band signals sampled at 20kHz. These signals were then down-sampled to 1,250Hz using the decimate function from the signal submodule of Scipy (version 1.11.2).

### Dentate spike detection

Dentate spikes were detected during sleep sessions from LFPs recorded from tetrodes located in the DG granule cell layer or silicon-probes with recording contacts in the DG granule cell layer. In silicon-probe recordings, we initially subtracted the LFP signals from all channels using a reference channel found in the stratum oriens. LFPs were band- pass filtered (1–200 Hz, using a 4th order Butterworth filter). The mean and standard deviation of the LFP amplitude were calculated across the entire sleep session and peaks that exceeded a threshold of six times the median absolute value of the filtered signals were designated as dentate spikes. The time bin with the largest local maximum was taken as the peak of the dentate spike, and this timestamp was recorded. If more than one peak appeared within a 50 ms frame, we retained only the highest amplitude peak. On recording days with several tetrodes in the DG, we used the tetrode with the largest mean DS amplitude to select DS event timestamps. Across all tetrode recordings we detected 32,215 DS events in total (mean ± SEM: 441.3 ± 29.2 per day, from 73 recording days in 12 mice); in silicon-probe recordings we detected 15,067 DS events in total (mean ± SEM: 1676.1. ± 316.5 per day, from 8 recording days in 3 mice).

### Sharp-wave ripple detection

For the LFPs of each pyramidal CA1 channel, we subtracted the mean across all channels (common average reference), band-pass filtered for the ripple band (80–250 Hz; 4th order Butterworth filter) and their envelopes (instantaneous amplitudes) were computed by means of the Hilbert transform. The peaks (local maxima) of the ripple band envelope signals above a threshold (5 times the median envelope of that channel) were regarded as candidate events. The onset and offset of each event were determined as the time points at which the ripple envelope decayed below half of the detection threshold. Candidate events passing the following criteria were determined as SWR events: (*i*) ripple band power in the event channel was at least twice the ripple band power in the common average reference (to eliminate common high frequency noise); (*ii*) each event had at least four ripple cycles (to eliminate events that were too brief); (*iii*) ripple band power was at least twice the supra-ripple band defined as 200-500 Hz (to eliminate high frequency noise, not spectrally compact at the ripple band, such as spike leakage artefacts). For events passing these criteria, the local maximum of each envelope was taken as the peak of the SWR, and these timestamps were recorded. On recording days with several tetrodes in the CA1 pyramidal layer, we used the tetrode with the largest mean ripple envelope amplitude to select SWR events. In tetrode recordings we detected 65,370 SWR events (mean ± SEM: 895.0 ± 82.3 per day, from 73 recording days in 12 mice).

### Peri-event time histograms (PETHs)

For analysis, we excluded all DS and SWR events that occurred within 50 ms of one another. We constructed PETHs over 400 ms windows, 200 ms either side of the peak DS amplitude or the peak of the SWR envelope, using a 1 ms bin width. The mean firing rate of each neuron was calculated during each 1-ms bin over the 400 ms window for each event. Z-scored firing rates were generated separately for each neuron by calculating the mean and standard deviation over the 400 ms PETH:

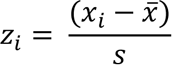

where z_i_ is the z-score at time bin *i*, *x_i_*is the firing rate in time bin *i*, *x̅* is the mean firing rate across all time bins, and s is the standard deviation of the firing rate across all time bins. The z-scored firing rate of each neuron was then smoothed using a 3-point moving average to eliminate spurious peaks in low firing rate neurons. For a cell to be classified as significantly activated during DS and/or SWR events, the firing rate within ± 20 ms of the event peak had to be > 3 standard deviations (s.d.) above baseline (calculated as the mean firing rate over the 400 ms window).

### Current source density analysis

Current sources and sinks were estimated from LFP recordings taken from single-shank 64-channel silicon-probes spanning the somato-dendritic axis of CA1 principal cells and reaching the inferior blade of the DG. LFP signals were first down- sampled to 1250 Hz. The current source density (Mitzdorf, 1985) unscaled signal at time *t* and electrode *n*, *CSD*[*t*]*n*, was estimated as:

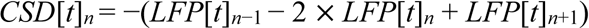

where *LFP*[*t*]*_n_*_−1_, *LFP*[*t*]*_n_* and *LFP*[*t*]*_n_*_+1_ are the LFP signals at time *t* recorded from neighboring electrodes (*n−1* and *n+1* are the channels immediately above and below *n,* respectively, with 20 μm spacing between electrodes). The silicon-probe recording site in the pyramidal layer was identified as the one with largest ripple-band power. We defined the location of radiatum and lacunosum moleculare layers according to the ripple and sharp-wave laminar profiles and electrode spacing (Lopes-dos-Santos *et al*., 2018). We sorted dentate spike events into Type 1

(DS_1_) or Type 2 (DS_2_) in the following way. First, we calculated the CSD estimates for all DSs at the peak of each event and used PCA to find the first two Principal Components from the resulting CSD traces. These principal components had as many dimensions as the number of silicon-probe channels (64). We then used a 2-component Gaussian Mixture Model to classify the events based on their projection onto the first two principal components. This consistently resulted in two event classes having the strongest sinks in different areas of the molecular layer. In line with previous research (Bragin *et al*., 1995; Dvorak *et al*., 2021), we classified the events with the strongest sink in the outermost part of the molecular layer as DS_1_, and events with their sink closer to the granular layer as DS_2_. Based on CSD classification, event proportions were DS_1_: 0.35; DS_2_: 0.65 (5274 DS_1_ versus 9793 DS_2_, based on 15,067 events from 8 recording days in 3 mice).

### Linear discriminant analysis classifier

To distinguish between DS_1_ and DS_2_ events using only the LFP traces, we trained a linear discriminant analysis (LDA) classifier using silicon-probe recorded LFPs from the granule cell layer (Castelli, 2023). LFP signals were first down-sampled to 1250 Hz and low-pass filtered at 50 Hz. We extracted 400 ms epochs centered around the peak of each DS (-200 to +200 ms, with 0.8 ms bin width), providing 500 time-based features (dimensions), one for each time bin, for each LFP trace. We then performed PCA on all silicon-probe-recorded DS LFP traces (15,067) to extract the number of components explaining 90% of the variance. This resulted in 16 principal components, which were then used to train a LDA classifier. We generated 20 models by, each time, randomly selecting 75% of the dataset, which was labelled as DS_1_ or DS_2_ based on the CSD classification described above, and then testing the classifier on the remaining (unlabeled) 25% of data. The classifier success rate was: median (IQR) = 85.4 (85.3-85.6) %. We then used the model with the highest accuracy to classify DS_1_ and DS_2_ events from LFPs recorded from the granule cell layer in our tetrode-recorded data.

From tetrode-recorded LFPs, the proportions of Type 1 and Type 2 DS events were: median (IQR) DS_1_ = 0.34 (0.25-0.38); DS_2_ = 0.66 (0.62-0.75), based on 10,337 DS_1_ versus 21,740 DS_2_ events in 73 recording days in 12 mice.

### Population spiking vectors

We generated event-based hippocampal population vectors of instantaneous principal cell spiking for every DS and SWR event using 50 ms wide windows centered on the peak of the DS or the peak envelope of the CA1 ripple (±25 ms from the peak). In addition, we calculated the spiking activity of hippocampal principal cells in equivalent 50 ms (‘no event’) control epochs, that contained neither DS nor SWRs. Baseline periods were selected from the same sleep sessions and excluded all epochs ± 250 ms either side of any DS or SWR events. To calculate the proportion of coactive neurons in each time window (**Figure 2G-H**), we calculated the number of simultaneously active hippocampal principal cells (i.e., cells firing at least one spike during the 50 ms window) by the total number of simultaneously recorded hippocampal principal cells. We then calculated the mean proportion of coactive cells for each recording session. For inclusion, each recording session required a minimum of 100 of each type of event (DS_1_, DS_2_, SWR) and a minimum of 20 simultaneously recorded hippocampal principal cells.

### Logistic regression classifier

We used a logistic regression classifier to distinguish between population vectors of hippocampal principal cell spiking activity during DS, SWR, or equivalent duration (50 ms) control vectors that were taken from 200–250 ms periods before the peak of either the DS or SWR events. For each recording session, we generated matrices of these population vectors (cells × epochs) for these four different event-types, and then binarized the spike counts (i.e., spike count > 0 = 1, else 0). For each recording day, we used the event with the lowest number of epochs to determine the training set size – for example, if there were 200

DS events, we used 75% (150 population vectors) as the DS training set, and randomly subsampled the SWR matrix for 150 SWR population vectors (with identical principal cells). This way, the training input to the classifier was balanced across event types. Similarly, the testing set consisted of the remaining (unlabeled) 25% of population vectors from the DS population vectors plus an equivalent number of SWR population vectors (e.g., 50 DS population vectors and 50 SWR population vectors, subsampled from the remaining SWR testing matrix).

For each recording day, we ran one model to classify event epochs, and another model to classify pre-event epochs. Model accuracy was measured as the proportion of correctly classified events (DS versus SWR, or pre-DS versus pre-SWR, respectively).

### Peer-to-peer coactivity analysis

We constructed hippocampal population graphs that represent the coactivity relationships between all pairs of principal cell spike trains recorded during a given sleep or exploratory session. These coactivity graphs were computed using 50 ms time windows for DS and SWR events and theta cycles as time windows for active exploratory sessions. To further control for the shared influence of the general network activity on peer-to-peer coactivity, we used for any two neurons (*i*, *j*) the regression coefficient *β* _*ij*_ obtained by fitting the GLM (**Figure 3A**):

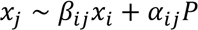

where *x*_*j*_, *x*_*i*_ are the z-scored event-nested spike trains of individual neurons *j* (the target) and *i* (the predictor), and *P* is the summed activity of the other *N* − 2 neurons,

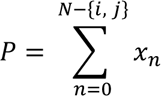

with *β*_*ij*_ weighting the influence of the population contribution to the activity of target neuron *j.* The recorded neurons (and their coactivity associations) are therefore the nodes (and their edges) in the coactivity graph of each task session. We described each graph by its adjacency matrix, *A*, as the *N* × *N* square matrix containing the pairwise coactivity relations within the network, yielding a weighted graph with no self-connections:

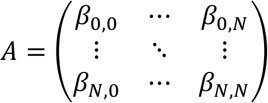

with *β*_*i*,*i*_ = 0 ∀*i in N*, and the symmetry in the weights of the network being ensured by setting 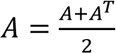 to form an undirected graph.

### Clustering coefficient

We computed a clustering coefficient to characterize the local synchronization of network activity by quantifying the number of three-node motifs. In each graph, for any neuron *i*, we obtained its clustering coefficient *C*_*i*_ using the formula proposed by Onnela et al. to quantify the strength of each triad (Onnela *et al*., 2005; Saramäki *et al*., 2007; Costantini and Perugini, 2014):

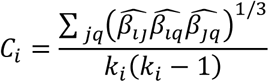

where *j* and *q* are neighbors of neuron *i*, all edge weights are normalized by the maximum edge weight in the network ^*β*^^ = *β*/ max(*β*), and *k_i_* is the degree of neuron *i*, which in these weighted graphs with no self-connection is equal to the number of neurons minus one. Note that this formula accounts for negative edges, yielding a negative value when there is an odd number due to the negative edges in the triad; it is positive otherwise. For this reason, this quantity can also be interpreted as a measure of the “structural balance” around a node, in the sense that its neighborhood presents coherent patterns of connectivity (Estrada, 2019).

### Population vector similarity

Population vectors of hippocampal principal cell spiking activity were generated for baseline, SWR, and DS events as described above, yielding separate (cell × event number) matrices of spike counts for each event-type. To remove potential biases caused by unequal numbers of events, we used the event-type with the fewest epochs to determine the final matrix size – for example, if there were 200 DS events in a given recording session, we randomly subsampled the SWR and baseline matrices to extract 200 SWR and 200 baseline population vectors (with identical principal cells) for comparison. Next we binarized these matrices (spike count > 0 = 1, else 0). Then we assessed the self-similarity for each event matrix (cells × event number) by computing the Pearson correlation coefficient for every pair of population vectors from the same event-type, and then calculating the mean across all of these correlation coefficients.

### Population dimensionality

We estimated the dimensionality of the principal cell population firing structure during SWRs and DSs from activity matrices that were matched for neuron identity and the number of DS and SWR events. We applied Principal Component Analysis (PCA) to each activity matrix, using the number of simultaneously recorded principal cells as the maximum number of components. Each matrix required at least 20 principal cells for inclusion in the analysis. We then extracted the number of components explaining 90% of the variance in these population vectors and scaled this by the total number of neurons in each matrix.

### Theta-cycle detection

Theta cycles were detected as described in Lopes dos Santos *et al*. (2023). Briefly, we used masked Empirical Mode Decomposition (Quinn *et al.,* 2021; https://pypi.org/project/emd/) to separate CA1 LFPs into oscillatory components termed intrinsic mode functions (IMFs). We delineated individual theta cycles from their troughs and peaks, i.e. the local maxima and minima of the theta IMF. Theta cycles were defined as peak-trough-peak sequences with trough-peak and peak-trough intervals between 31-100 ms and peak-to-peak distances between 71-200 ms.

### Reactivation of waking coactivity patterns

We leveraged our pairwise peer-to-peer coactivity measure (as described above and in **Figure 3A**) to estimate DS and SWR reactivation. We compared the tendency of principal cell pairs to co-fire in theta cycles during exploration (theta coactivity) with the tendency to co-fire in DS or SWR during the following sleep/rest period (post-DS or post-SWR co-firing), controlling for their baseline co-firing in the sleep/rest period before (pre-DS or pre-SWR co-firing). From the resulting GLM, we extracted the β coefficients of the linear regression that predicted post-SWR or post-DS coactivity from theta coactivity, controlling for pre-SWR or pre-DS coactivity, respectively. We tested the significance of these β coefficients in two ways. First, we performed control GLMs using the pre-DS (or pre-SWR) coactivity as the dependent variable and the theta coactivity and post-DS (or post-SWR) coactivity as the independent variables. Second, we constructed a random probability distribution of β weights for theta coactivity by shuffling the cell pair identity, thereby generating a null distribution (based on 1000 GLMs, each time randomly shuffling cell-pair identity).

### Closed-loop optogenetic intervention

Real time detection of DSs was achieved by first high pass filtering the DG LFP signals (5 Hz) using the on-board signal processing capabilities of the Intan RHD evaluation board (RHD2000, Intan Technologies, USA) and triggering a laser pulse if the LFP signal exceeded a voltage-threshold. Thresholds for DS-onset detection were set for each mouse during a sleep session at the start of each recording day so that DS events were consistently detected (∼3 S.D. above mean signal amplitude). Threshold detection triggered a digital transistor-transistor logic (TTL) output pulse from the RHD interface to a Master 8 stimulation timing device (A.M.P.I., Jerusalem, Israel), which in turn sent a 100 ms duration square-wave pulse to activate the laser. In the ‘DS-synchronized’ condition, the laser was triggered with zero latency from DS-onset, whereas in the ‘DS-delay’ condition the laser was triggered 100 ms after DS detection (**Figure 5A**). The laser delivered yellow-green light (561- nm) into the dentate gyrus, which in Grm2^DG^::ArchT mice activated the outward proton pump, Archaerhodopsin T to suppress DG granule cells. To investigate changes in firing rates in individual hippocampal principal cells during light-delivery, we constructed PETHs over 400 ms windows, 200 ms either side of DS-onset, using a 1 ms bin width, and then z-scored these binned spike trains. To quantify light firing responses, we extracted the maximum z-score after DS- onset (**Figure 5C**) and the summed z-score (**Figure S5B**) between DS-onset and 100 ms after DS-onset for each hippocampal principal cell.

### Novel object recognition memory task

On each task day, mice explored a square-walled open field (**Figure 5D**; the ‘object arena’) containing four objects, each positioned midway along a given wall, ∼1-cm from the wall edge. During the first session in the object arena, mice explored four completely novel objects (‘sampling’ session, 10 min). After the sampling session, mice were placed into a sleep box where they received DG-targeting light delivery that was either synchronized to DS-onset (DS-synchronized condition) or delayed by 100 ms from DS-onset (DS-delay condition), as described above (sleep/rest session, 20 min). Before the start of the next test session, one of the four objects was replaced with a different (and completely novel) object, and mice then explored the four objects again (‘test 1’ session, 10 min). This process was repeated, with another sleep session (∼20 min, with either DS-sync or DS-delay light-delivery), followed by another object exploration session with one completely novel object and three previously encountered objects (‘test 2’ session, 10 min). During each test session, we measured the time spent exploring each object and we calculated the percentage time spent investigating the novel object versus the (mean) percentage time spent investigating the familiar objects (i.e., those objects seen in the previous session). On any given day, mice received the same light-delivery condition (i.e. both sleep sessions were either DS-sync or DS-delay). There was no difference in the number of DS-onset events detected and therefore the number of light pulses delivered across the two conditions (mean ± SEM: DS-synchronized = 351 ± 81; delay = 377 ± 83; paired permutation test: p = 0.7) and there was no difference in the number of SWR events detected across the two conditions (mean ± SEM: DS-synchronized = 570 ± 98; delay = 393 ± 128; paired permutation test: p = 0.4).

### Tissue processing and immunohistochemistry

At the completion of experiments, mice were deeply anesthetized with pentobarbital and perfused transcardially with 0.1 M PBS followed by 4% paraformaldehyde (PFA) in PBS. Brains were extracted and kept in 4% PFA for ∼24–72 h and then transferred to PBS (with 0.05% sodium-azide). For immunostaining, free-floating sections (50-μm) were rinsed in PBS with 0.25% Triton X-100 (PBS-T) and were blocked for 1 hour at ∼20 °C in PBS-T with 10% normal donkey serum (NDS). Sections were then incubated with primary antibodies diluted in 3% NDS blocking solution and incubated at 4 °C for 72 hours (GFP anti-chicken, 1:1,000, Aves Labs, catalog no. GFP-1020; NeuN guinea pig, 1:500, Synaptic Systems, catalog no. 266 004). All sections were rinsed three times for 15 min in PBS-T and incubated for 4 hours at ∼20 °C in secondary antibodies in the blocking solution (Cy3 donkey anti-guinea pig, 1:400, Jackson ImmunoResearch, catalog no. 706-165-148; goat anti-chicken 488, 1:1,000, Thermo Fisher Scientific, catalog no. A-11039). Sections were then rinsed three times for 15 min in PBS-T, with some sections then incubated for 1 min with DAPI (0.5 μg ml^−1^, Sigma, D8417) diluted in PBS to label cell nuclei before three additional rinse steps of 10 min each in PBS. Sections were mounted on slides, cover-slipped with Vectashield (Vector Laboratories, catalog no. H-1000) and stored at 4 °C. Sections were also used for anatomical verification of the tetrode tracks. Images were acquired using a Zeiss confocal microscope (LSM 880 Indimo, Axio Imager 2) with a Plan-Apochromat ×20/0.8 M27 objective and the ZEN (Zeiss Black 2.3) software.

### Statistical analysis

Analyses were performed in Python 3.8 (https://www.python.org/downloads/release/python-3816/), using the Python packages DABEST (Ho *et al*., 2019), scipy (Virtanen *et al*., 2020), numpy (Harris *et al*., 2020), matplotlib (Hunter, 2007), seaborn (Waskom, 2021), pandas (McKinney, 2010), scikit-learn (Pedregosa *et al*., 2011), statsmodels (Seabold and Perktold, 2010). Error bars, mean ± S.E.M unless otherwise stated. We used throughout this study a bootstrap-coupled estimation of effect sizes, plotting the data against a mean difference between the left-most condition and one or more conditions on the right and compare this difference against zero using 5,000 bootstrapped resamples. In these estimation graphics (DABEST plots), each black dot indicates a mean difference and the associated black ticks depict error bars representing 95% confidence intervals; the shaded area represents the bootstrapped sampling-error distribution. Bandwidth estimates for the kernel density estimate were computed using the scikit-learn package. All statistical tests were performed two-sided, unless otherwise stated. Paired permutation tests (or equivalent paired tests) were performed for repeated-measures analyses and unpaired tests used for independent samples.

## Data availability

The datasets generated during and/or analyzed during the current study will be made available via the MRC BNDU Data Sharing Platform (https://data.mrc.ox.ac.uk/) on reasonable request.

## Code availability

The software used for data acquisition and analysis are available using the web links mentioned in the methods. Code for the DS_1_ / DS_2_ classifier is available at https://github.com/mcastelli98/DentateSpikeClassifier.

## Acknowledgements

We would like to thank B. Micklem for technical assistance; all members of the Dupret lab for feedback during the project. This work was supported by the Biotechnology and Biological Sciences Research Council UK (Awards BB/S007741/1, BB/N002547/1) and the Medical Research Council UK (Programmes MC_UU_12024/3 and MC_UU_00003/4).

## Author contributions

Conceptualization, S.B.Mc. and D.D.; Methodology, S.B.Mc., V.L.d.S., M.C., G.P.G., S.T. and D.D.; Formal Analysis, S.B.Mc.; Investigation, S.B.Mc. and S.K.E.T.; Resources, S.B.Mc., V.L.d.S., G.P.G., K.H. and D.D.; Writing – Original Draft, S.B.Mc. and D.D.; Writing –Reviewing & Editing, S.B.Mc, V.L.d.S, G.P.G., M.C., K.H., S.K.E.T., and D.D.; Visualization, S.B.Mc. and D.D.; Supervision, D.D; Funding Acquisition, S.B.Mc. and D.D.

---

**Figure S1.**
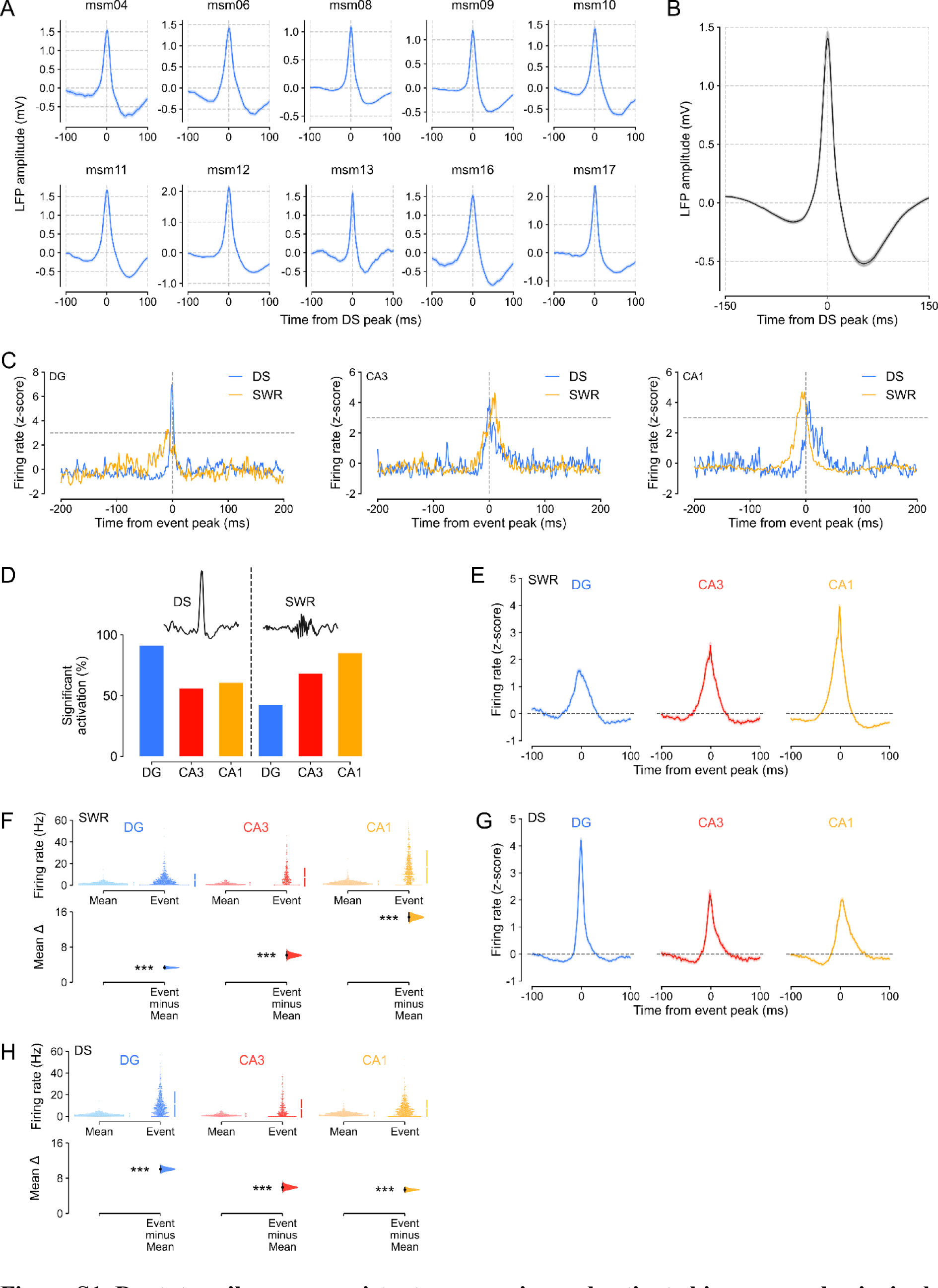
**Dentate spikes are consistent across mice and activate hippocampal principal cells.** **(A)** Examples of mean ± SEM dentate spike (DS) LFP waveforms from individual mice. **(B)** Group mean ± SEM DS LFP waveform (n=73 sessions, 12 mice). **(C)** Examples of three individual principal cells’ z-scored firing rates during DSs (blue) and SWRs (orange). Cells were considered significantly activated if the z-score > 3 (within ± 20 ms of the event peak, shown at time 0). The three example cells were significantly activated by both events. **(D)** Percentage of significantly activated principal cells, as defined in (C), during DS (left) and SWR (right) events. **(E-H)** Peri-event time histograms (E,G) showing z-scored firing rates ± 100 ms around the event peak and estimation plots (F,H) comparing the overall mean firing rate of each principal cell (calculated across the entire recording session) to its peri-event firing rate (calculated as the mean firing rate ± 5 ms around the peak of the event) during SWRs (E,F) and DSs (G,H). \*\*\**P* < 0.001.

**Figure S2.**
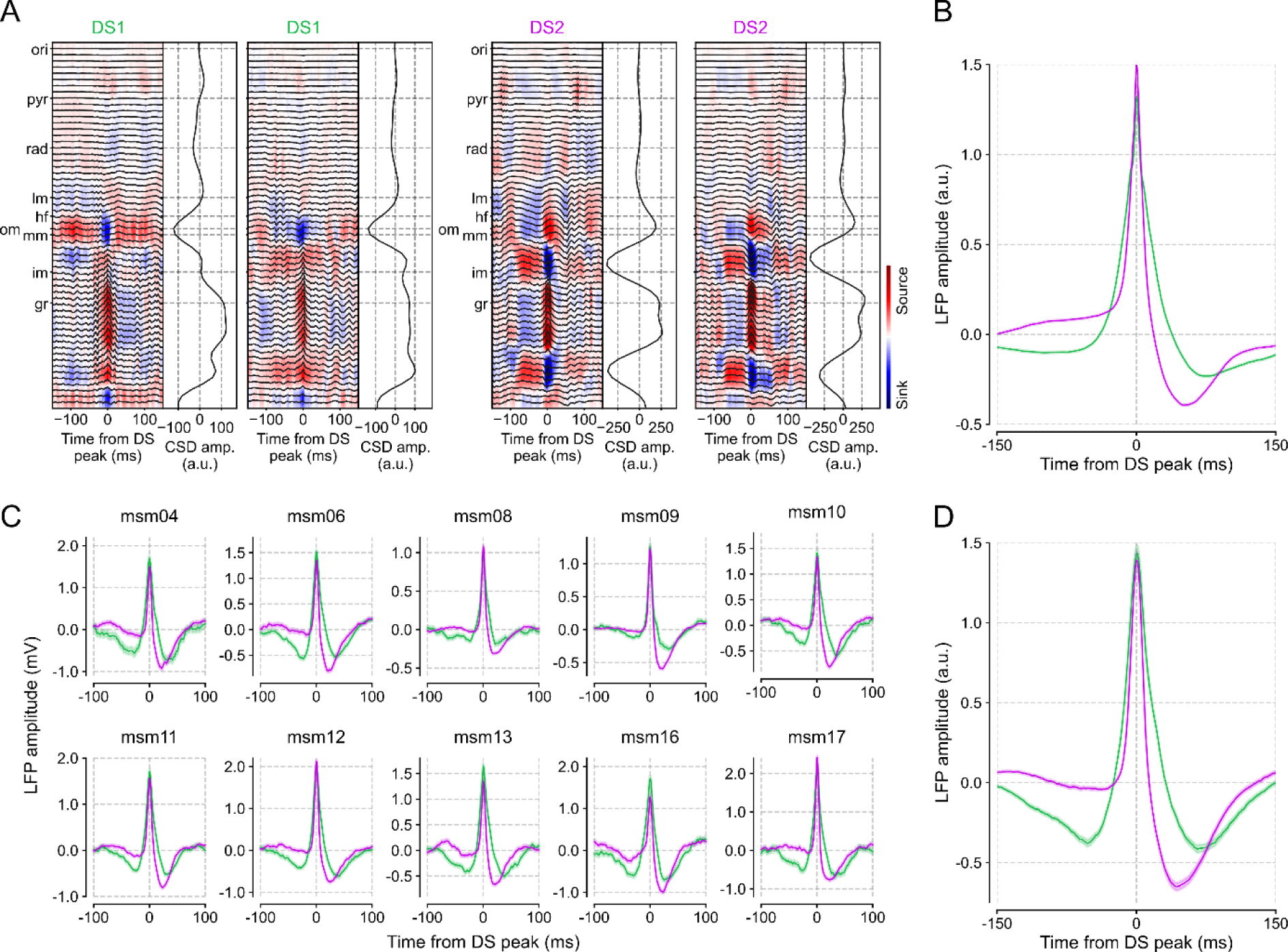
Current source density and local field potential profiles for DS_1_ and DS_2_ events. (A) Examples of current source density (CSD) profiles for individual Type 1 (DS_1_) and Type 2 (DS_2_) dentate spikes (two DS_1_ examples, two DS_2_ examples) recorded from a 64-channel silicon-probe. *Left panel*: the instantaneous CSD ±150 ms around the event peak. *Right panel:* the CSD amplitude at each depth (based on the mean amplitude from –25 to + 25 ms around the event peak). Stratum oriens: ori; pyramidal layer: pyr; stratum radiatum: rad; lacunosum moleculare: lm; hippocampal fissure: hf; outer molecular layer: om; middle molecular layer: mm; inner molecular layer: im; granule-cell layer: gcl. (B) Group mean ± SEM LFP waveforms for DS_1_ and DS_2_ events from silicon-probe recordings (8 sessions, 3 mice). (C) Examples of mean ± SEM LFP waveforms for DS_1_ and DS_2_ events from tetrode recordings in individual mice. (D) Group mean ± SEM LFP waveforms for DS1 and DS2 events from tetrode recordings (73 sessions, 12 mice).

**Figure S3.**
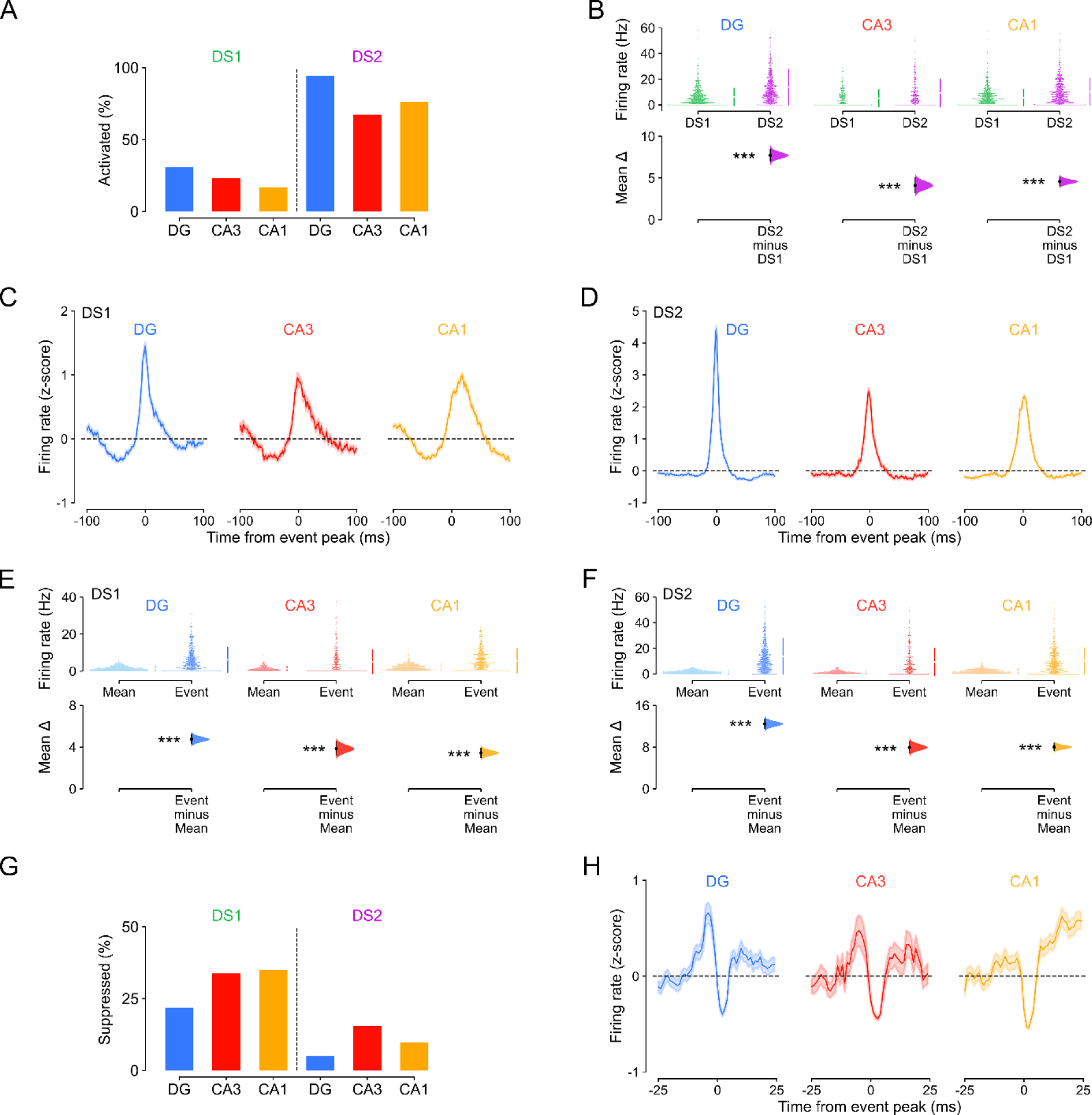
DS_1_ and DS_2_ events increase the average hippocampal principal cell spiking. **(A)** Percentage of significantly activated principal cells (i.e., cells with firing rate > 3 s.d. above baseline ± 20 ms either side of event peak) during DS_1_ and DS_2_ events. **(B)** Comparison of firing rates during DS_1_ versus DS_2_ events (mean rate during ± 5 ms around the event peak; DG n= 921, CA3 n=388, CA1 n= 887 principal cells, in 12 mice). **(C-F)** Peri-event time histograms (C,D) showing z-scored firing rates ± 100 ms around the event peak and estimation plots (E,F) comparing overall mean firing rates (calculated across the entire recording session) to peri-event firing rate (calculated as the mean firing rate ± 5 ms around the peak of the event) for all principal cells during DS_1_ and DS_2_ events. DG n=921, CA3 n=388, CA1 n=887 principal cells (12 mice). **(G)** Percentage of suppressed principal cells (i.e., cells with a z-score < 0 during the event peak) during DS_1_ and DS_2_ events. **(H)** Peri-event time histograms showing z-scored firing rates ± 25 ms around the event peak for the lowest quartile of activated principal cells (i.e., the 25% least activated / suppressed principal cells) during DS_1_ events. (DG n=230, CA3 n=97, CA1 n=221 principal cells). \*\*\**P* < 0.001.

**Figure S4.**
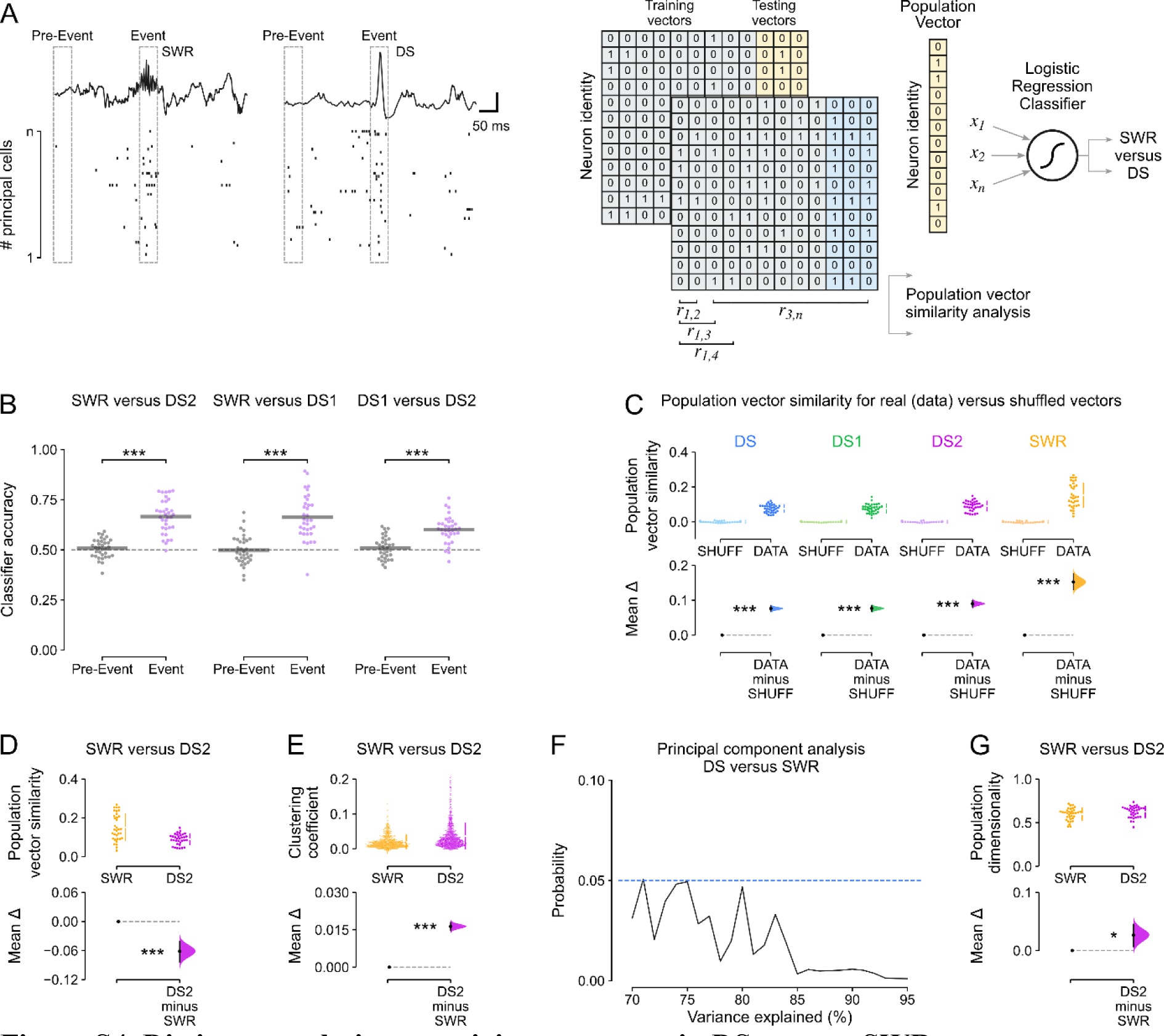
Distinct population coactivity structures in DSs versus SWRs. **(A)** Toy example illustrating construction of cell × event spiking activity matrices. Logistic regression classifiers were trained using 75% of the population spiking activity vectors and tested with the remaining 25% of vectors, on any given session. We used the event type with the lowest number of epochs to determine the training and testing set size and then randomly subsampled the other event matrix to generate the same number of training and testing vectors for each event type, so that the classifier was balanced across event types. Separate matrices and classifiers were utilized for event epochs and pre-event epochs. **(B)** Classifier performance for SWR versus DS_2_, SWR versus DS_1_, and DS_1_ versus DS_2_ population vectors. In these analyses, the inclusion criterion was that the session had to contain a minimum of 20 principal cells and 100 events of each type. **(C)** Population vector similarity for all event types compared to their control condition in which each event vector was correlated with ‘shuffled’ population spiking vectors, where the cell identity was randomly shuffled. **(D)** Population vector similarity was higher for SWR than DS_2_ events. **(F)** PCA to compare the dimensionality of SWR versus DS matrices (cell × event number), matching the number of events for each event type, determining the number of components required to explain 70–95% of the variance. In each case, the dimensionality was significantly higher for DS versus SWR events at α < 0.05 (Wilcoxon test for paired samples, 1-tailed). **(G)** Population dimensionality was higher for DS_2_ than SWR events. *\*P* < 0.05, \*\*\**P* < 0.001.

**Figure S5.**
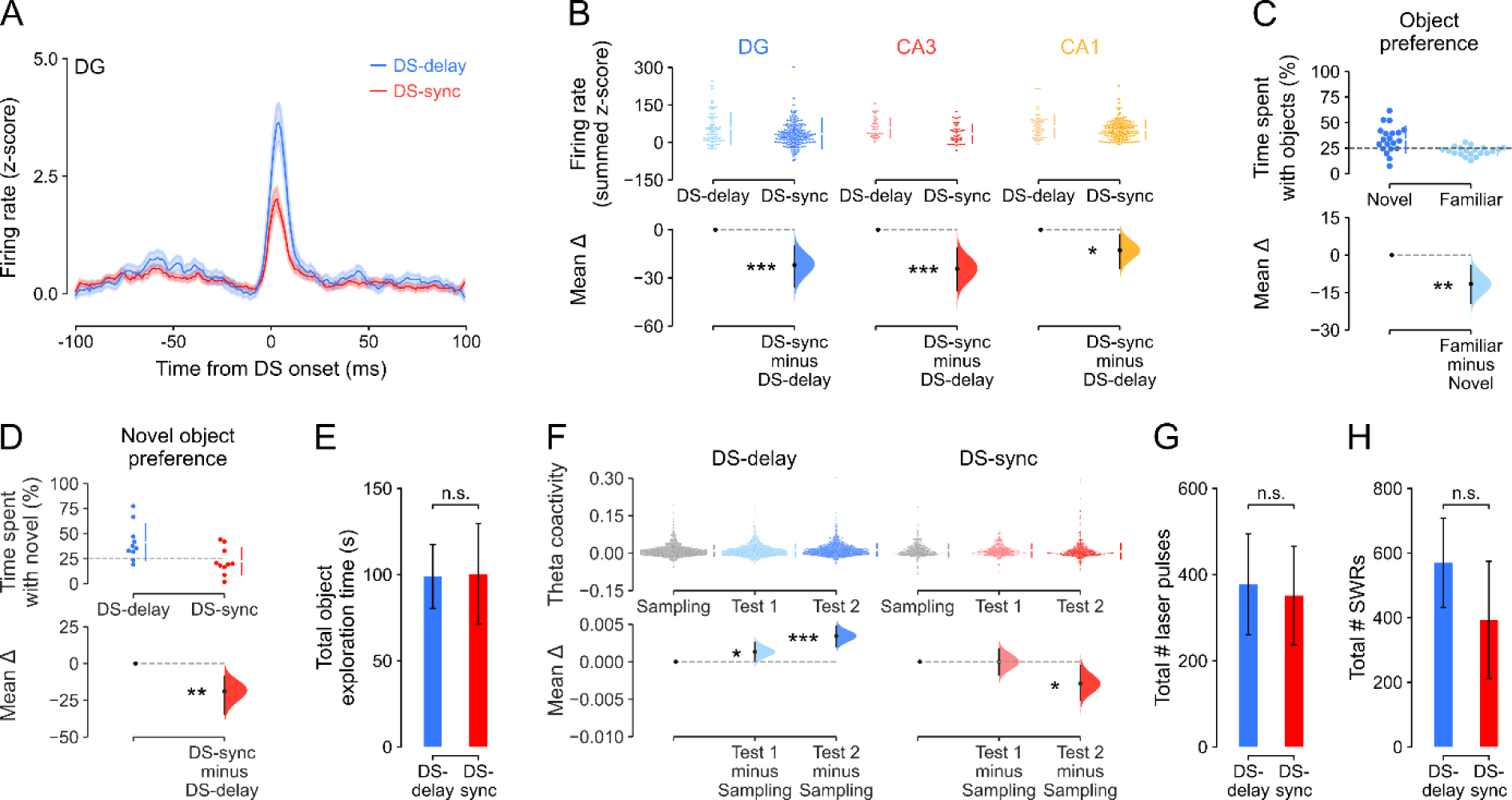
Optogenetic suppression of dentate granule cells during DSs reduces CA spiking, impairs novel object recognition memory and associated theta coactivity patterns. **(A)** Firing response of dentate granule cells (DGCs) to DSs in Grm2^DG^::ArchT mice, either with light-delivery (to inhibit DGCs) synchronized to DS-onset (DS-sync, red line) or with light-delivery offset by 100 ms after DS-onset (DS-delay, blue line; Delay: n=127; DS: n=319 DGCs in 5 mice). **(B)** The summed z-scored firing rate was significantly lower in the DS-sync than DS-delay condition for hippocampal principal cells. Estimation plot showing the effect size for the summed z-scored firing rate (0–100 ms after DS-onset) in DGCs (blue), CA3 (red) and CA1 (orange) during DSs in the Delay versus DS-sync conditions. Upper: each data point represents the summed z-scored firing rate for each neuron. Lower: mean difference in the summed z-score between the two conditions (DGC: Delay: n=127; DS: n=319; CA3: Delay: n=58; DS: n=70; CA1: Delay: n=98; DS: n=276, in 5 mice). **(C)** Estimation plot for object preference in a ‘no-laser’ control group of mice, showing significantly more time spent investigating the novel object (n=20 test sessions in 5 mice). **(D)** Estimation plot showing that novel object preference was significantly higher in the DS- delay than the DS-sync condition (n=10 test sessions per condition, in 3 mice). **(E)** Total time spent exploring the objects did not differ between the DS-delay and DS-sync conditions. **(F)** Theta peer-to-peer coactivity during the novel object recognition task increased over the three sessions of object exploration in the DS-delay condition but decreased in the DS-sync condition. Paired estimation plot showing theta coactivity during Test 1 and Test 2 relative to the initial novel object exploration (Sampling). **(G)** The number of laser pulses delivered did not differ between the DS-delay and DS-sync conditions. **(H)** The number of SWRs detected during sleep/rest did not differ between the DS-delay and DS-sync conditions. \**P* < 0.05, \*\**P* < 0.01, \*\*\**P* < 0.001.

**Supplementary Table 1.** Generalized linear model analysis of SWR and DS reactivation of waking theta coactivity patterns.

| Event | Dependent variable | Independent variables | Df | F | Prob. | R<sup>2</sup> |
| --- | --- | --- | --- | --- | --- | --- |
| SWR | Post-exploration | Theta coactivity<br>Pre-exploration | 2 (model),<br>7007 (resid.) | 971 | P < 0.0001 | 21.7% |
| SWR | Pre-exploration | Theta coactivity<br>Post-exploration | 2 (model),<br>7007 (resid.) | 890 | P < 0.0001 | 20.2% |
| DS | Post-exploration | Theta coactivity<br>Pre-exploration | 2 (model),<br>7007 (resid.) | 635 | P < 0.0001 | 15.3% |
| DS | Pre-exploration | Theta coactivity<br>Post-exploration | 2 (model),<br>7007 (resid.) | 585 | P < 0.0001 | 14.3% |

| Event | Dependent variable | Independent variables | $\beta$ coefficient | CI (95%) | t | Prob. |
| --- | --- | --- | --- | --- | --- | --- |
| SWR | Post-exploration | Theta coactivity | 0.214 | [0.186, 0.242] | 14.8 | P < 0.0001 |
|  |  | Pre-exploration | 0.438 | [0.415, 0.461] | 37.6 | P < 0.0001 |
| SWR | Pre-exploration | Theta coactivity | 0.128 | [0.101, 0.155] | 9.4 | P < 0.0001 |
|  |  | Post-exploration | 0.384 | [0.364, 0.404] | 37.6 | P < 0.0001 |
| DS | Post-exploration | Theta coactivity | 0.347 | [0.285, 0.408] | 11.1 | P < 0.0001 |
|  |  | Pre-exploration | 0.341 | [0.320, 0.362] | 32.1 | P < 0.0001 |
| DS | Pre-exploration | Theta coactivity | 0.203 | [0.138, 0.268] | 6.2 | P < 0.0001 |
|  |  | Post-exploration | 0.376 | [0.353, 0.399] | 32.1 | P < 0.0001 |

